# Stochastic Modeling of Biophysical Responses to Perturbation

**DOI:** 10.1101/2024.07.04.602131

**Authors:** Tara Chari, Gennady Gorin, Lior Pachter

**Affiliations:** Division of Biology and Biological Engineering, California Institute of Technology, Pasadena, California; Fauna Bio, Emeryville, California; Department of Computing and Mathematical Sciences, California Institute of Technology, Pasadena, California

## Abstract

Recent advances in high-throughput, multi-condition experiments allow for genome-wide investigation of how perturbations affect transcription and translation in the cell across multiple biological entities or modalities, from chromatin and mRNA information to protein production and spatial morphology. This presents an unprecedented opportunity to unravel how the processes of DNA and RNA regulation direct cell fate determination and disease response. Most methods designed for analyzing large-scale perturbation data focus on the observational outcomes, e.g., expression; however, many potential transcriptional mechanisms, such as transcriptional bursting or splicing dynamics, can underlie these complex and noisy observations. In this analysis, we demonstrate how a stochastic biophysical modeling approach to interpreting high-throughout perturbation data enables deeper investigation of the ‘how’ behind such molecular measurements. Our approach takes advantage of modalities already present in data produced with current technologies, such as nascent and mature mRNA measurements, to illuminate transcriptional dynamics induced by perturbation, predict kinetic behaviors in new perturbation settings, and uncover novel populations of cells with distinct kinetic responses to perturbation.

## Introduction

Over the past decade, advances in high-throughput sequencing technologies and techniques for multi-condition, multi-sample experimentation have enabled genome-wide perturbation experiments that can assay hundreds of genetic interventions or drug combinations simultaneously [1–4]. The effects of these perturbations can be measured at single-cell resolution across multiple measurement types (or ‘modalities’) [5, 6]. The goal of such work is to unravel how the processes of DNA/RNA regulation produce observed cellular responses and diversity. This provides fertile ground for mechanistic studies of transcription, to tease apart how features such as DNA structure, genetic interactions, and mRNA processing dynamics [6–8] regulate processes like cellular differentiation and disease progression. For example, in many published single-cell RNA sequencing (scRNA-seq) datasets [9], one can obtain information about nascent and mature (‘unspliced’ and ‘spliced’) mRNA expression, which represents different components of mRNA production and processing [10]. Such information is particularly relevant for understanding mRNA splicing dynamics which play important roles in normal development as well as cancer progression and resistance acquisition [11–13].

However most tools for analysis of large-scale perturbation datasets focus on observational effects only, e.g., changes in expression, using only mature mRNA information, rather than modeling the generative, transcriptional processes themselves [1, 4, 14, 15]. Deep learning approaches also focus on prediction of expression patterns [14, 16], but even when considering multiple modalities [5], the physical interpretation of the learned parameters and relationships between the measurements can be hard to extract. These tools and approaches also often require several transformations of the data, to remove noise or enhance biological signal, which can themselves incur distortion and obscure interpretation [17–19]. Mechanistic approaches and investigations are often limited to smaller or more homogeneous systems [20–22], or assess modalities and their corresponding dynamics, independently [23].

In this work we demonstrate how the extension of stochastic models of transcription to noisy, high-throughput perturbation datasets alternatively enables perturbation analyses through the underlying, biophysical processes of DNA/RNA regulation. Using the unspliced and spliced count modalities, we can uncover condition-specific kinetics, predict regulation of transcription kinetics in combined perturbation settings, and define novel cell states induced by perturbation (Fig. 1). Thus, we can generate hypotheses about *how* perturbations affect the transcription and processing of RNA, for downstream investigation and experimentation.

**Figure 1:**
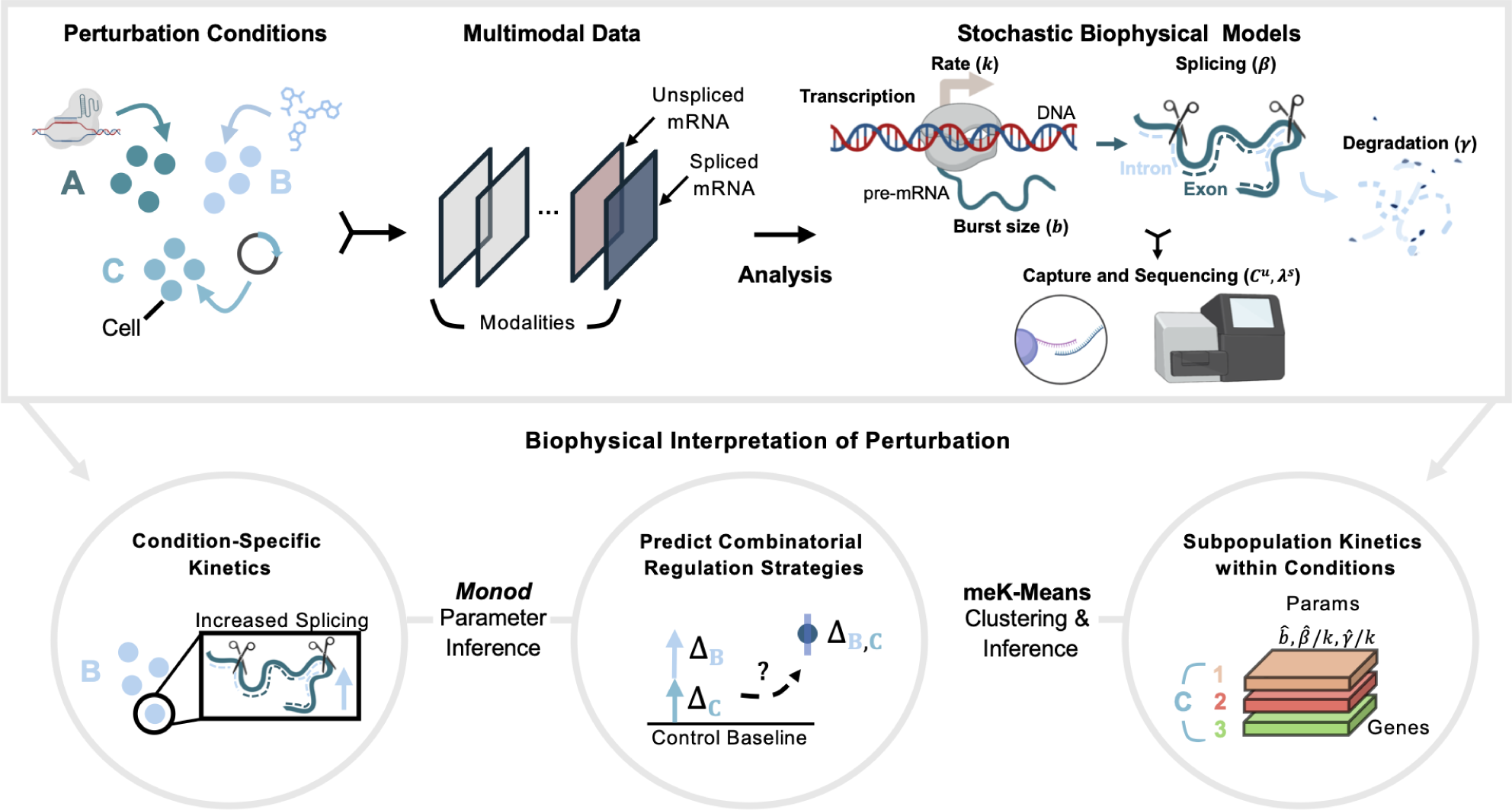
Stochastic Models for Biophysical Interpretation of Perturbation Data. (Top Box) From left to right, generation of perturbation data where different perturbants are applied to groups of cells. Data is processed into count matrices, which can then be analyzed through stochastic, biophysical models. (Bottom Circles) Biophysical analyses of perturbation data that can be performed using the *Monod* package for CME-based inference and the meK-Means clustering algorithm. Created with BioRender.com.

## Results

For our analysis, we take as input scRNA-seq perturbation datasets with unspliced and spliced gene count matrices, which represent nascent and mature mRNA molecule counts [9, 24]. Such counts can be obtained by realignment of existing data to an intron-annotated reference using tools such as kallisto [9, 10, 24, 25]. The datasets in our analysis encompass both drug-based perturbations, A549 lung cancer cells under Dexamethasone (DEX) treatment [26] and mouse neural stem cells (NSCs) under 96 different drug combinations [3], as well as genetic intervention assays, specifically (single and dual) CRISPRa [2] and CRISPRi [27] perturbations in K562 cells (leukemia cell line) (see Supplementary Table 1).

These data represent discrete, sparse, and noisy molecular count information, where the unspliced and spliced counts represent non-independent entities, related through the transcriptional processes of the central dogma. Thus we use a Chemical Master Equation (CME)-based formalism to model these count data and the biophysical processes they derive from (see Methods) [28, 29]. Briefly, the biological (and technical) variation of a gene’s molecule counts are modeled through the transcription of the gene at rate *k* and burst size *b* (corresponding to a ‘bursty’ model of transcription [29, 30]). The gene’s mRNA molecules are then spliced at a rate *β*, and degraded at a rate *γ*. The length-biased, poly(A) capture of the mRNA molecules used in the technologies of the above datasets, and subsequent molecule sampling over the sequencing pipeline, are represented through modality-specific capture parameters *C^u^, λ^s^* (with *u, s* corresponding to unspliced and spliced modalities). At steady-state, the rate of transcription is not uniquely identifiable, thus throughout this analysis we report relative splicing and degradation rates *β/k, γ/k*, relative to the transcription rate [28].

With the *Monod* package [31] for parameter inference of CME models from single-cell data, we can then fit the biological (and technical) parameters which define this biophysical model of the sequencing data, for the individual conditions in these datasets (see Methods). This produces gene-specific parameters for each perturbation condition of interest. We denote the inferred estimates of parameter *θ* as *θ*^^^, i.e., ^^^*b, β*^^^*/k, γ*^*/k*.

However, this analysis relies on external annotations of the groups of cells to be assessed, as gene parameters are inferred over the distribution of all cells in the annotated condition. Yet it is often of interest to cluster data in an unsupervised manner to discover heterogeneous perturbation responses and cellular states potentially *within* a perturbation condition [4, 32]. For this we use the meK-Means (‘mechanistic K-Means’) algorithm described in [33], which we have incorporated into the *Monod* framework, to combine clustering of scRNA-seq data with biophysical inference. The meK-Means method simultaneously learns populations of cells present in a heterogeneous dataset alongside the transcriptional kinetics that define those populations, providing a natural mechanism for integration of modalities for the common analysis task of clustering.

With these tools, we then obtain from datasets (1) inferences of changes in transcription kinetics induced by perturbations, (2) predictions of kinetics in dual perturbation conditions, from single-perturbation condition parameters, and (3) identification of cell populations displaying distinct perturbation responses. With perturbation-specific biophysical parameters, as in (1), we can define DE-*θ* genes as genes displaying ‘differential’ behavior. For the ‘DE-*θ*’ analyses below, we consider log_2_ fold changes (FCs) greater than 2, at the parameter level (for a given parameter *θ*) between two conditions, e.g., the perturbed condition and a control condition. This is a more general notion of differential expression (DE), as compared to the standard definition in scRNA-seq analysis, based on differences in mean spliced expression [34].

Likewise, we extend the definition of a ‘marker’ gene based on the parameter-level DE. Interpretation of a marker at the parameter level is highly context dependent; if one is looking for increased mRNA stability between two populations, lower degradation would denote a marker gene of one population, whereas if higher turnover is of interest, higher degradation would classify the gene as a marker for the other population. For the purposes of this analysis, we denote a gene as a marker when burst size is increased or both splicing and degradation are decreased (i.e., increased burst frequency or transcription rate *k*). If neither is the case, an increase in splicing or decrease in degradation (increased mRNA stability) denotes a marker.

To predict and assess the effect of perturbation on kinetic parameters in dual-perturbation conditions, as in (2), we use additive and multiplicative models of the perturbation’s effects given the corresponding, individual perturbation conditions (see Methods). Effectively, these models assume that the changes to transcription, i.e., burst size or transcription rate, in the single-perturbation conditions (as compared to the control condition) are additive or multiplicative in nature in the combined perturbation condition (see Methods).

For simultaneous clustering and parameter inference, as in (3), we then apply meK-Means under the same length-biased model of the sequencing data used to infer condition-specific parameters. MeK-Means takes as input the unspliced and spliced matrices as well as a user-defined number of clusters, K, though the algorithm can converge to fewer numbers of clusters. This produces cluster and gene-specific biophysical parameters. Thus, given the gene counts of cells in a perturbed setting, we can learn populations of cells demonstrating similar responses and their defining, kinetic parameters. The same DE-*θ* analysis as described above, can then be applied to these learned parameters.

### Kinetic Effects of Perturbation on Transcription

To better understand the *how* behind a perturbation’s effect on gene expression, e.g., mRNA production, we used *Monod* to fit the biophysical parameters of the model in Fig. 1,5 on 3,000 genes (which contained a minimum number of unspliced and spliced counts) for the DEX-treated A549 cells at 0 hours of treatment and 2 hours of treatment (see Methods). The single-cell indexing and labeling technique sci-fate was used to generate these data [26]. For each condition (Fig. 2a), we extracted the burst size, splicing, and degradation parameters across genes (Fig. 2b) (see Methods). All unspliced and spliced mRNA counts, which combine the 4sU labeled and unlabeled counts captured in this experiment, were used for parameter inference. As described in the original study [26], we found that the degradation rates between the two conditions displayed high correlations (Fig. 2b). However, we could additionally assess these correlations relative to the correlations between burst sizes and splicing rates of the two conditions, highlighting greater differences in these parameters across conditions, as opposed to degradation rates (Fig. 2b). All three parameters were also fit under the same model of transcription, as opposed to fitting separate kinetic models for the parameter(s) of interest [26].

**Figure 2:**
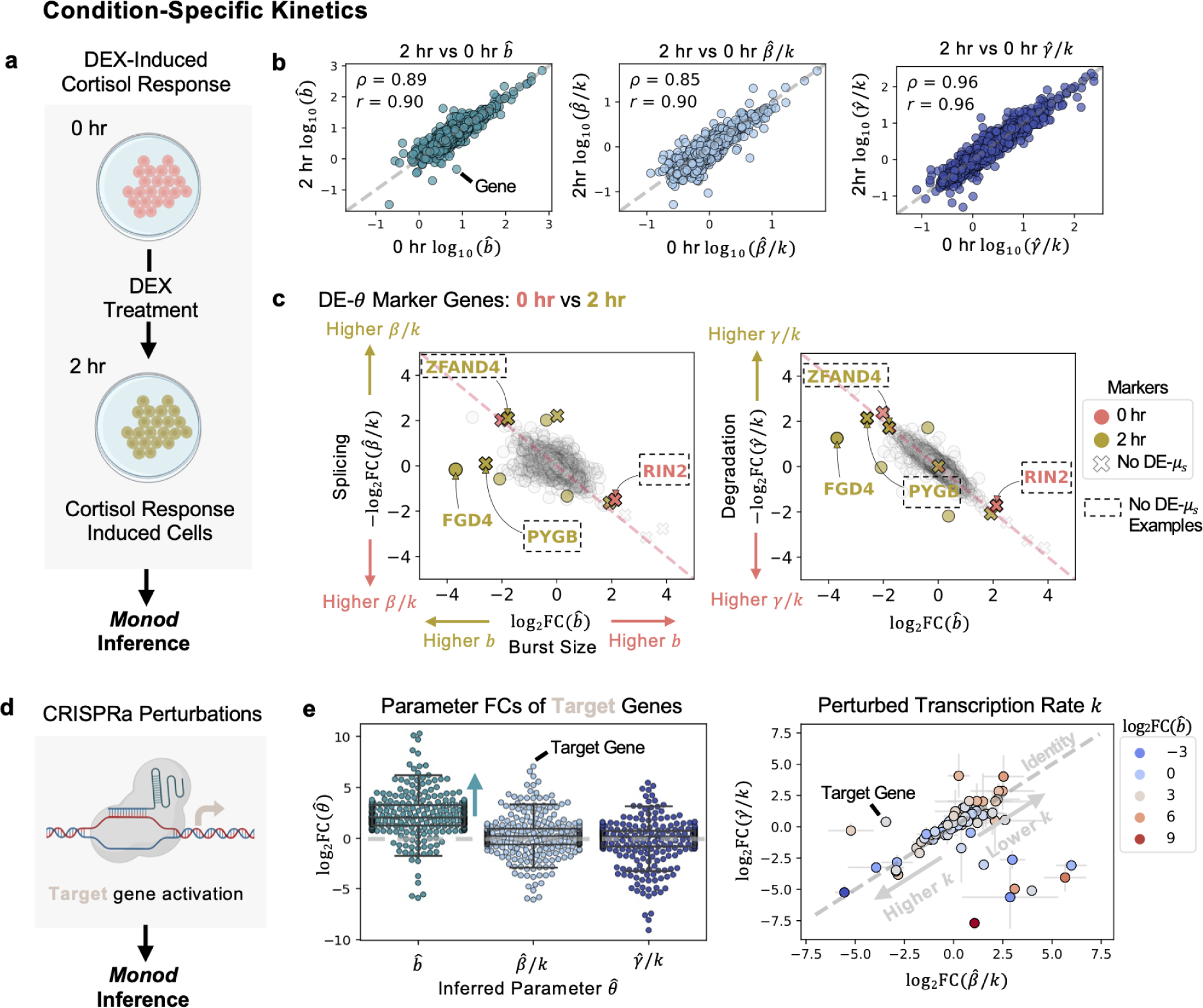
Perturbation Condition-Specific Kinetics. **a** Diagram of DEX-treated A549 cells, at 0 and 2 hours. Parameter inference for data done using *Monod*. **b** Inferred parameters, for burst size, splicing, and degradation compared between cells at 0 and 2 hours of treatment. Spearman and Pearson correlation are denoted by *ρ* and *r* respectively. **c** ‘DE’ or ‘Differentially-Expressed’ genes at the parameter level (*θ*) shown between the clusters corresponding to 0 and 2 hour°cells. Genes in dashed box denote genes where DE is detected at the parameter level but not at the observed, mean S-level. (Left) Splicing rate vs burst size shown for genes. (Right) Degradation rate versus burst size shown. Grey genes denote ambiguous markers, or non-significant FCs. **d** Diagram of CRISPRa perturbation used for *Monod* parameter inference. **e** (Left) fold change (FC) of inferred parameters for the target (activated) genes. FCs shown as compared to all controls in the study. (Right) FCs for degradation versus splicing parameters for the target genes. Error bars denote standard deviation of FC across all controls. Genes shaded by burst size FC. Created with BioRender.com.

The original study noted a difficulty in using whole transcriptome information (labeled and unlabeled transcripts) to assess differences in the cells between the 0 hour and 2 hour condition [26]. It appeared that standard highly variable gene (HVG) selection and dimensionality reduction of the data resulted in an inability to separate the 0 hour and 2 hour responses, without solely focusing on the newly transcribed (labeled) mRNA or using a procedure to combine the individual principal components (PCs) of the labeled and unlabeled data after PCA reduction. However, by fully describing the joint distribution of the unspliced and spliced counts through biophysical parameters, we have the ability to not only differentiate the perturbation responses of the 0 hour and 2 hour conditions through DE-*θ* genes, but also detect these differences for genes that do not necessarily show discernible FCs when comparing mean expression between conditions, i.e., that are not DE-*µ_s_*, as is done in classical DE analyses (Fig. 2c). For example, we can detect changes in the burst size of the cortisol response marker *FGD4* [26, 35] as well as the changes in burst size and splicing or degradation, for the gene *ZFAND4* (a prognostic and metastasis marker) [36], the glycogen phosphorylase gene *PYGB* [37] induced in other stress conditions, and the *RIN2* GTPase gene involved in endocytosis and membrane trafficking [38], which did not display FCs at the mean spliced expression-level (Fig. 2c). We additionally separate the two treatment conditions through the biophysical clustering approach, below (see Uncovering Perturbed Populations with Distinct Kinetics).

To further validate and explore the kinetic realizations of *Monod* inference in various perturbation settings, we fit biophysical parameters for each intervention condition (with greater than 50 cells) in the CRISPRa (activation) Perturb-seq study [2], which encompassed single and dual gRNA conditions (Fig. 2d). For this dataset, samples were generated with the 10x Genomics v2 protocol. The CRISPRa mechanism increases transcriptional output by potentially recruiting and stabilizing components of the transcription preinitiation complex [39]. In each activation condition, we found that burst sizes of the corresponding target genes were increased (with, on average, log_2_FCs greater than 2) (Fig. 2e left), mirroring burst size observations discussed in the Narta activation protocol, which recruited artificial transcription factors to the transcription site [40]. In comparison, average splicing and degradation rate FCs were near zero (Fig. 2e left). However, in cases where both the splicing and degradation rate FCs are in the same direction (sign), we can infer potential changes to the denominator of these relative rates, i.e., in the transcription rate *k* (Fig. 2e right) [31]. This reveals genes where transcription rate, or burst frequency, is altered in addition to, or in spite of, changes in burst size, suggesting different strategies for the gene’s transcriptional regulation (Fig. 2e right) [41–43].

Beyond the target genes themselves, we can ask if other genes were activated in a similar manner, such as through increased burst size, and investigate what genomic features might produce these shared kinetic behaviors. As an example, we looked for genes with increased burst sizes (log_2_FC *>* 2) in the dual activation condition where *CBL* and *CNN1* were activated (leading to differentiation of the K562 cells). We found genes associated with erythroid differentiation, such as *ALAS2* [44], and transcription factors such as *FOXA1* and *EGR1*. By looking at genomic positions of all such genes, we then assessed if shared localization may contribute to these kinetic similarities (Supplementary Fig. 1a). Using nuclear speckle proximity scores from a lymphoblastoid cell line in [45] (see Methods), we found that genes with increased burst sizes, particularly those clustered on chromosome 19, were potentially in speckle ‘close’ genomic regions. Nuclear speckles represent nuclear bodies that are hubs for increased pol II density and transcriptional activity (as well as splicing machinery), and regions demonstrating inter-chromosomal interactions (Supplementary Fig. 1b) [45]. Thus, through the inferred biophysical parameters, we could identify genes with similar bursting behavior and develop hypotheses about how their physical organization could lead to this property.

The statistical framework of our inference model additionally allows for assessment of uncertainty in these inferred parameter estimates and their information content or how much information (as defined through the Fisher Information Matrix (FIM)) the data carries about that parameter (see Methods). We then investigated how properties of the intervention conditions tested, such as the number of cells sampled or the UMIs detected per cell, lead to more or less informative estimates of the biophysical parameters of interest in those conditions (Supplementary Fig. 2). The strongest relationship was between average UMIs per cell and the information content of the parameter inferred, where higher UMI detection (akin to ‘read-depth’) resulted in more informative parameter estimates, potentially plateauing at around an average of 8.5-9k UMIs per cell, for an average of 1,900 genes detected (Supplementary Fig. 2). This information-based analysis provides a physically-informed route for followup experimental design [20].

By taking advantage of the full joint distribution between the modalities in these datasets, we can not only quantitatively assess the kinetic effects of a perturbation, but also detect changes in these kinetic parameters not discernible in conventional, expression-based analyses. The added resolution afforded by kinetic inferences allows for the development of more focused hypotheses for follow-up experimentation.

### Predictive Models of Combinatorial Perturbations

Many methods for analysis of perturbation data additionally focus on predicting the effects of perturbations in novel settings [14, 16, 46]. This can aid in simulation of responses, and minimizing experimental efforts for downstream investigations. Often these predicted changes or effects are defined as changes in spliced expression, a proxy for changes in transcription. However, different underlying mechanisms may contribute to these observed changes. Previous work has described potential models of how perturbations or regulatory inputs in combination can impact transcription kinetics [47], and in turn downstream expression [23, 47], but application of such models is not generally extended to single-cell genomics perturbation data.

Additionally, it is non-trivial to use predicted spliced and unspliced counts to predict underlying, mechanistic effects. If tools predict changes in mean expression [1], this does not provide enough distributional information to infer dynamics. When counts are modeled more explicitly, the distributions parametrizing the observed counts from multiple modalities are independent, ignoring causal relationships and making physical interpretation of the learned parameters difficult [5, 14].

However, by using the inferred parameters from *Monod* in single-perturbation conditions, we can define models at the level of the kinetic parameters to predict the parameters in dual conditions (i.e., where both perturbations are present). The parameters of the single-perturbations then inherently describe their full joint distributions of unspliced and spliced count, and extend this description to the dual-perturbation setting. This also extends previous investigations, which assess the additive and multiplicative properties of mean spliced expression under perturbation [23], to the behavior of the governing rates which produce those observed behaviors.

Given the inferred parameters in the single-guide conditions in the CRISPRa Perturb-seq study, and the inferred parameters in the control conditions, we can test the ability of simple models of additive and multiplicative behavior to recapitulate the kinetic parameters in dual-guide conditions (see Methods). Specifically, we assessed how well multiplicative and additive models of the changes in burst size and transcription rate describe the observed changes in parameters in the combined conditions (Fig. 3a) (see Methods).

**Figure 3:**
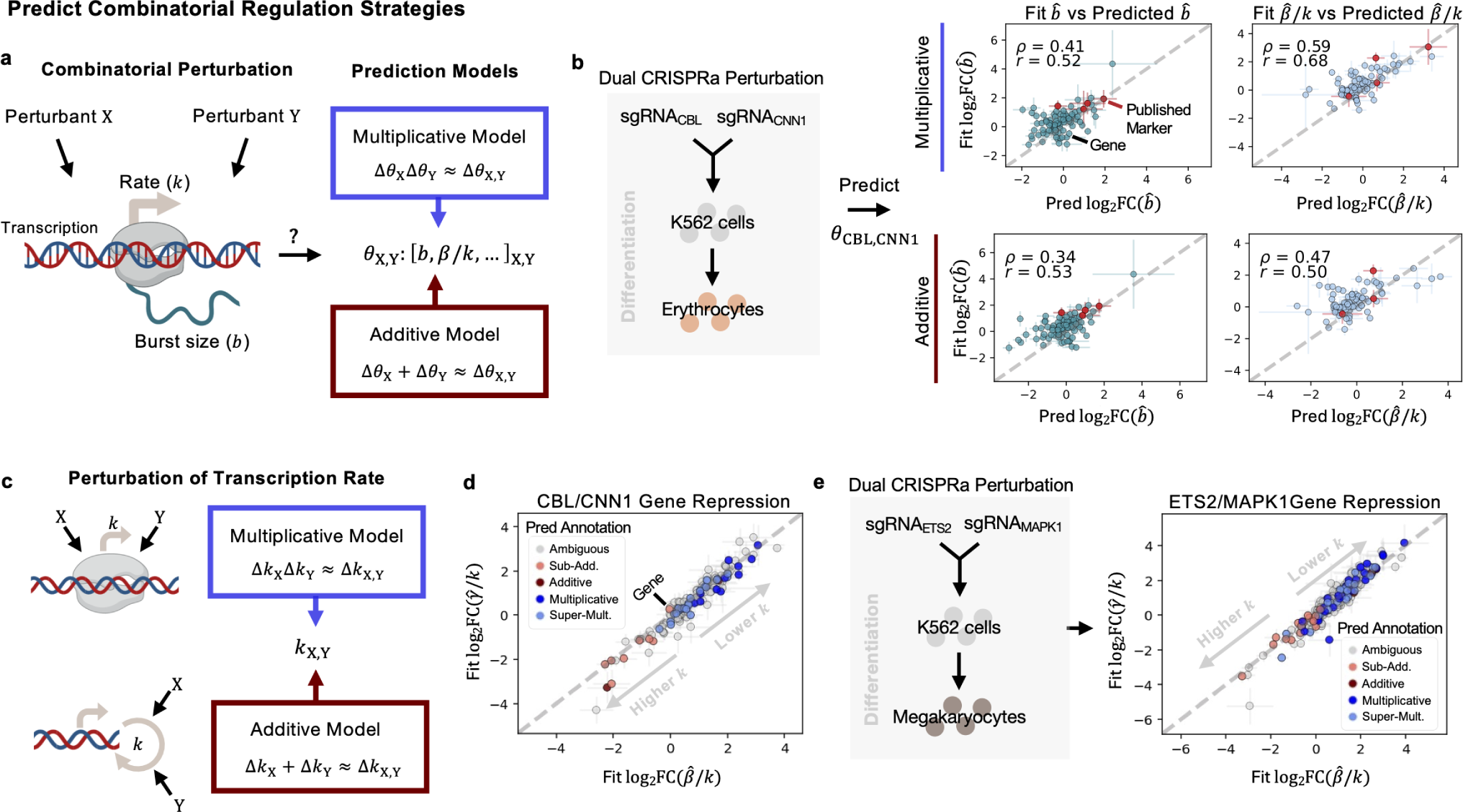
Prediction of Combinatorial Regulation Strategies. **a** Diagram of potential effects of perturbants on transcription. Multiplicative and additive prediction models predict parameters in combined perturbation conditions from single perturbation conditions. **b**Predicted parameters for the dual CRISPRa condition, for genes *CBL* and *CNN1*. Predictions from both models shown for burst size and splicing, correlated against the ‘Fit’ FC, from the inferred parameters in the combined condition. Spearman and Pearson correlation are denoted by *ρ* and *r* respectively. **c** Diagram of potential models of perturbant effects on transcription rate *k*, in a single-step transcription model. **d** FCs of the inferred degradation rate versus splicing rate, in the combined *CBL/CNN1* condition. Error bars denote standard deviation of FC across all controls. Genes colored by the best-fitting predictive model. **e** FCs of the inferred degradation rate versus splicing rate, in the combined *ETS2/MAPK1* condition. Error bars denote standard deviation of FC across all controls. Genes colored by the best-fitting predictive model. Created with BioRender.com.

The genes tested in these predictive models were selected in a similar fashion to the genes selected for analysis in the original study, where a random forest regression model was used to select genes that separated well the single-guide and dual-guide conditions from the control condition [2]. However, we did not include the dual condition in regression-based selection of genes, as we treat this condition as unseen. For these genes, we see positive correlations, *>* 0.5, for at least one models’ predictions as compared to the observed FCs for the inferred (‘Fit’) parameters, across the dual conditions described in the [2] (Supplementary Fig. 3a, Fig. 3b). Fig. 3b displays the correlation across all genes tested, of the predicted burst size and relative splicing rates in the dual condition, described above, where genes *CBL* and *CNN1* were activated. Genes in red denote ‘Published Marker’ or genes described in the original text as markers of the combined condition [2]. Interestingly, the more heterogeneous dual condition [2], with *DUSP9* and *MAPK1* activated, was not as well-described by both predictive models (Supplementary Fig. 3a). We then tested these predictive models on other data types, including dual CRISPRi conditions [27] and dual drug conditions (where low/mid/high ranges of concentrations of EGF, BMP4, or retinoic acid, RA, were added to NSCs) [3] (Supplementary Fig. 3a). We found similarly positive correlations between the predicted parameter changes and observed changes, and could distinguish when, for example the additive model better described changes in burst size than the multiplicative model, and vice versa (Supplementary Fig. 3a). The positive correlations of the predicted parameters were also higher than the correlations produced by a negative control model (Supplementary Fig. 3b), where parameters from single, control guide conditions replaced the single-guide conditions in the predictive models (representing how well random noise could predict the fit parameter changes). Generally, these selected genes which better separated the single-guide (perturbed) cells from the control, performed better under the predictive models than looking across all genes (Supplementary Fig. 3c).

Interpreting what these additive or multiplicative changes in kinetic parameters mean in terms of transcriptional regulation strategies being employed by the cell can be difficult, particularly if changes induced by the individual conditions are in different directions. Thus we focused interpretation on scenarios where the transcription rate *k* was likely affected (splicing and degradation rate FCs were in the same direction/of the same sign in both single-conditions). Given our bursty transcription model’s assumption of the single forward rate *k*, additive effects on the transcription rate in the combined condition can be framed as deriving from parallel and independent effects of individual interventions to catalyze the reaction rate, *k* [47] (see Methods). In the multiplicative case, additive effects of the interventions that affect recruitment and binding energy (e.g., of RNA pol II or other transcriptional activators) combine to produce multiplicative effects at the level of the rate *k* [47] (see Methods) (Fig. 3c).

We then delved into two dual-guide conditions from the CRISPRa Perturb-seq study, where the individual guides demonstrated transcriptional effects in the same direction, likely inducing differentiation towards erythrocyte (Fig. 3d) or megakaryocyte (Fig. 3e) lineages [2], and selected for repressed genes, i.e., where there was a negative FC in unspliced counts (see Methods). We assigned the most representative predictive model to the observed changes in the parameters of the dual-conditions (see Methods). Though for many genes it was ambiguous whether the additive or multiplicative model fit better [23], among the genes where we we could discern more model-specific behavior, we found a dominance of the multiplicative and super-multiplicative predictions when the transcription rate was lowered (Fig. 3d,e). This included genes such as *ATF5* ( in the *CBL*/*CNN1* condition) whose downregulation has been noted in neuronal differentiation [48], and *SNHG29* in the *ETS2* /*MAPK1* condition, a non-coding RNA which can repress the effects of miR-210-3p [49], which in turn positively regulates K562 differentiation [50]. For such genes, this suggests use of a more recruitment-based strategy as described above, potentially with non-independent effects of the interventions [47], to enact repression.

To further explore potentially shared mechanisms behind these repressed genes (Supplementary Fig. 1a), we looked for enrichment of transcription factor binding sites in their promoter regions using the HOMER package for motif discovery [51] (see Methods). For genes demonstrating multiplicative and super-multiplicative repression of their transcription rates, we found that over half had ETS binding domains, particularly for the Elk-1 transcription factor which has repressive domains that can act cooperatively to dampen activity of activation domains (in *cis* and in *trans*) [52], demonstrating a possible shared regulatory strategy.

Even with simpler models of how perturbations act in combination, by approaching the prediction task from the level of the kinetic parameters, we can expand previous works assaying multiplicative and additive behaviors at the expression-level, to the kinetics which underlie these observations and provide hypotheses of regulation strategies employed by the cell.

### Uncovering Perturbed Populations with Distinct Kinetics

In addition to prediction of kinetic parameters across modalities, such as unspliced and spliced counts, it is a non-trivial problem to discover and define populations of cells demonstrating distinct perturbation responses given multiple molecular measurements. Most standard approaches to clustering scRNA-seq data do not use multiple modalities at once [4, 53], use heuristics to map between clusters or neighborhoods if given individual modalities [54], or use non-physically-interpretable deep learning methods to integrate the modalities for clustering [55]. This then results in multiple, potentially arbitrary choices for a user to make when deciding how to combine the modality-specific matrices for clustering or determining which method’s clustering results to proceed with [33]. To this end, we applied the meK-Means clustering algorithm to simultaneously learn populations or clusters of cells in heterogeneous perturbation conditions, where ‘clusters’ are defined as cells displaying similar transcriptional kinetics, i.e., the processes of DNA/RNA regulation we are interested in investigating with perturbation (Fig. 4,6) [33].

**Figure 4:**
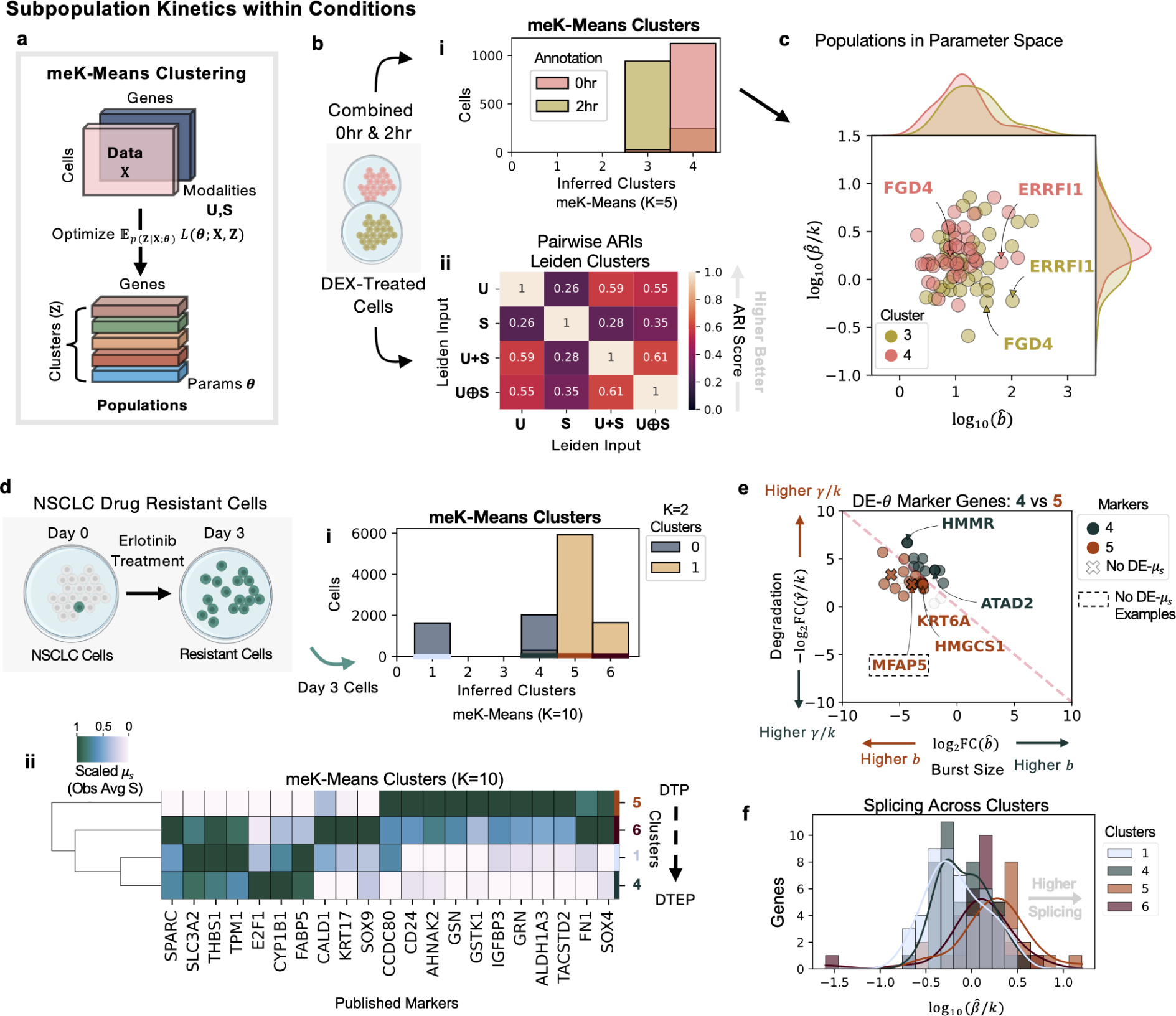
Inference of Subpopulation Kinetics within Conditions. **a** Diagram of the meK-Means algorithm for clustering **b** Diagram of combined 0 and 2 hour DEX-treated cells, then passed to meK-Means or Leiden for clustering. **i** Barplot of cluster assignments from meK-Means shown, where K=5, and distribution of 0 and 2 hours cells between them. **ii** Pairwise ARI scores between the Leiden clustering results given various input matrix options. **c** Plot of inferred splicing versus burst size parameters for the inferred clusters 3 and 4. Density plots of the respective parameter distributions shown per cluster (top and side of plot). **d** Diagram of drug resistant NSCLCs after 3 days of erlotinib treatment. Day 3 cells passed to meK-Means for clustering. **i** Barplot of cluster assignments from meK-Means, where K=10, and the distribution of the two populations of cells from the meK-Means K=2 clustering between them. **ii** Hierarchical dendrogram plot of meK-Means inferred clusters based on mean expression (scaled across columns) of published marker genes. **e** ‘DE’ or ‘Differentially-Expressed’ genes at the parameter level (*θ*) shown between the inferred clusters 4 and 5. Genes in dashed box denote genes where DE is detected at the parameter level but not at the observed, mean S-level. Degradation rate versus burst size shown. Grey genes denote ambiguous markers, or non-significant FCs. **f** Histograms of splicing rates shown for all genes across the inferred clusters. Created with BioRender.com.

**Figure 5:**
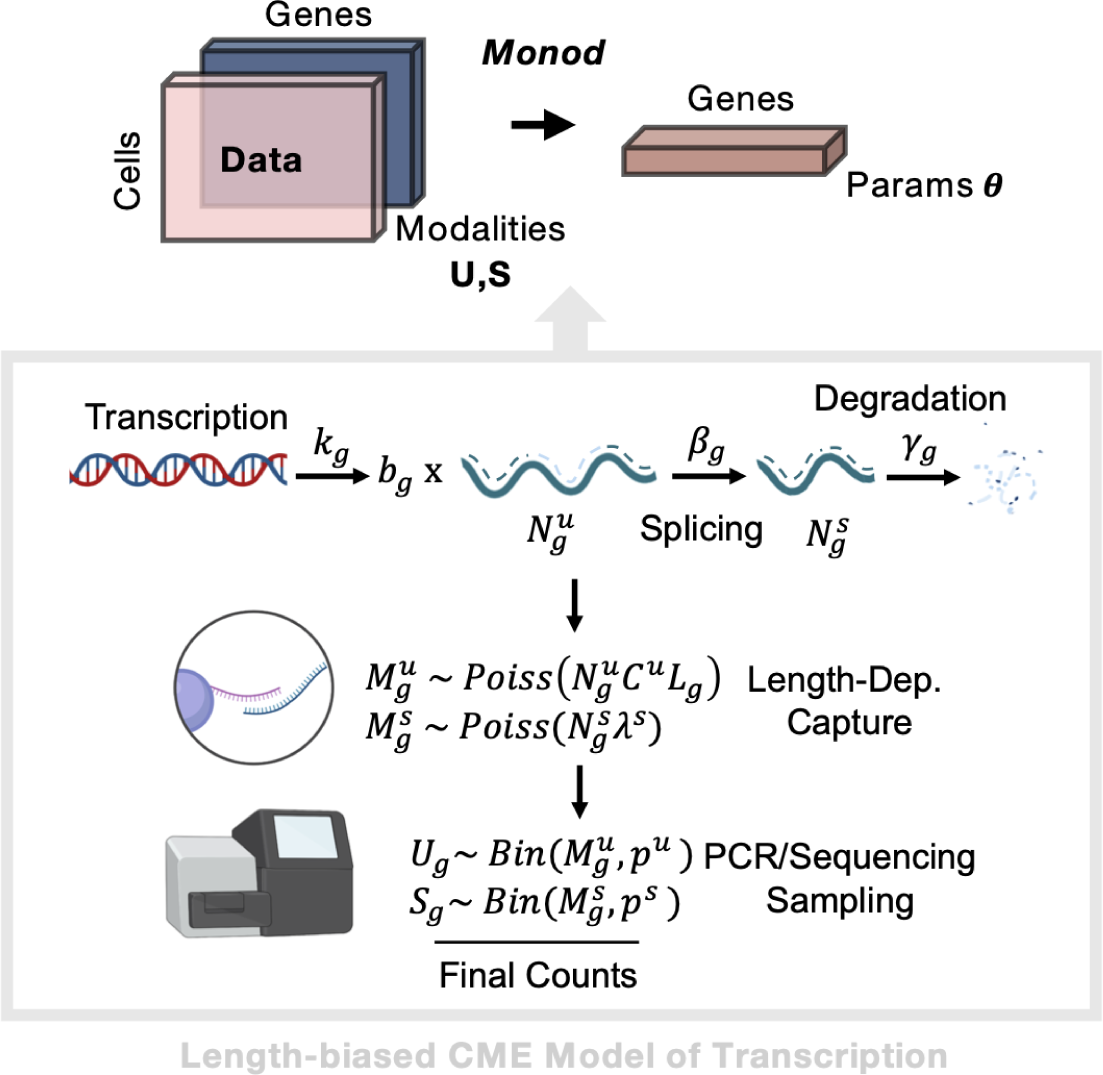
Parameter Inference for Length-Biased CME Model. (Top) Workflow from input data matrics, through *Monod* inference, to output biophysical parameters per gene. (Bottom) Description of the steady-state CME Model for bursty transcription and length-biased capture of molecules during sequencing. Created with BioRender.com.

**Figure 6:**
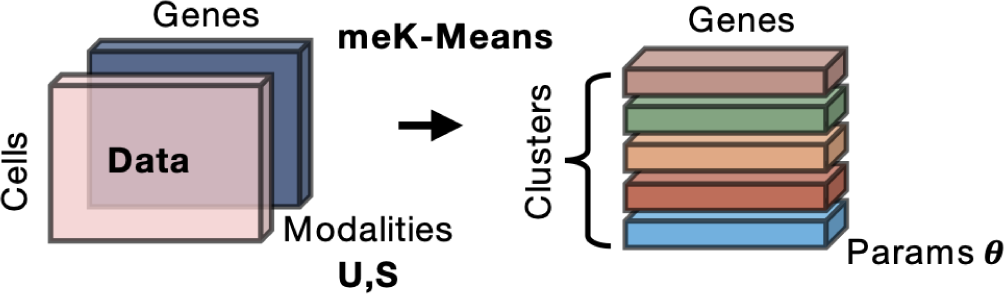
meK-Means Inference for Clustering. Workflow from input data matrics, through meK-Means inference, to output biophysical parameters per gene and per learned cluster.

We first demonstrated the use of meK-Means on the DEX-treated A549 cells, combining cells from 0 and 2 hours of treatment (Fig. 4b). As described in the original study [26], it was difficult to separate these two populations using whole transcriptome information and standard scRNA-seq processing pipelines. However, given a previously published list of genes potentially implicated in the cortisol response [35], and filtered for sufficient unspliced and spliced counts as well as overdispersed behavior (see Methods) [33], meK-Means was able to clearly separate the treated cells (Fig. 4b i). Given that only 45 genes remained after filtering, this demonstrates the importance of quality over quantity in gene selection, and potential pitfalls of standard HVG selection in cases like this, or other more ad hoc procedures. Additionally, even with the same genes, standard Leiden clustering [53] for example, leaves users with multiple, discordant cluster assignments of the same data, since multiple possible input matrices could be used, without biophysical rationale for using one over the other (Fig. 4b ii, Supplementary Fig. 4a). Here **U** + **S** represents the summation of the individual **U** and **S** matrices, and **U** *⊕* **S** represents the concatenation of the **U** and **S** matrices, or a more independent treatment of the modalities. The Adjusted Rand Index (ARI) scores, calculated between all pairs of clustering results, represents similarity between the cluster assignments, where 1.0 means overlapping assignments, and 0.0 means non-overlapping or random assignments. Leiden clustering, among the other possible clustering techniques offered in packages such as Seurat [54] and scanpy [56], can also be combined with a host of integration and embedding techniques for multimodal data [57, 58] leading to more ad hoc choices for a user employ, without physically-interpretable results.

Effectively, meK-Means clusters cells in biophysical parameter space, as shown in Fig. 4c, where parameters of genes with validated transcriptional changes during the cortisol response (e.g., *FGD4* and *ERRFI1*) [26, 35] stand out from each other between the two populations. The meK-Means-inferred parameters for these genes also corresponded near-identically to the parameters previously inferred from the conditions separately (described above in Kinetic Effects of Perturbation on Transcription) (Supplementary Fig. 4c).

We then applied meK-Means to cells without clear treatment partitions, where PC9 cells, an EGFR-mutant non-small cell lung cancer (NSCLC) cell line, were treated with erlotinib, a common first-line treatment for NSCLC, and sequenced after three days (following the 10x Genomics v3 protocol) [59]. At Day 3, multiple drug-resistant populations of cells had developed [59]. In the development and persistance of drug-resistant cancer cells, the kinetics of splicing as well as transcription are particularly relevant to how these cells acquire resistance and proliferate [12]. The Day 3 cells were thus clustered with meK-Means, using genes from both ‘classical’ HVG selection [56] and genes from the literature potentially marking resistance development, again filtered for overdispersed behavior and minimum unspliced and spliced counts (Fig. 4d). From this, we found four clusters of cells (Fig. 4d i), belonging to two larger populations of cells as also described in the original study (Supplementary Fig. 5c) [59]. These four populations spanned drug-tolerant persister (DTP) and drug-tolerant expanded persister (DTEP) states described in the study [59], which can persist and proliferate in the presence of drug treatment. For example, populations representing more DTP-like states demonstrated greater expression of the resistance marker *TACSTD2*, while DTEP-like states expressed the marker *CYP1B1* at higher-levels (Fig. 4d ii, Supplementary Fig. 6, Figure 1g in [59]).

We then extracted DE-*θ* genes between the inferred populations, specifically more DTP or more DTEP clusters, e.g., 4 and 5. Markers included the microfibril-associated gene *MFAP5*, which did not display high FCs at the mean spliced-level, but did display differing burst size, splicing, and degradation rates between the populations (Fig. 4e). DE-*θ* genes also included genes with differential behavior between the DTP and DTEP states in the original study, such as downregulation of *KRT6A* (associated with epithelial development) [59] and *HMGCS1* (associated with cholesterol metabolism) [59] in the DTEP cluster (4) (Fig. 4e). We additionally found reduced degradation of the *HMMR* gene between the populations, a prognostic marker gene in several other human cancers [60] (Fig. 4e). Since splicing dynamics in resistant populations are also of interest, we examined the distribution of splicing rates in each population. This revealed increased splicing rates overall in the DTP-like populations as compared to the DTEP population (Fig. 4f), suggesting potentially more aberrant splicing behavior in these populations [12].

This biophysical approach to clustering perturbation data, brings together the count modalities under a self-consistent model of the biology, and highlights not only which cells demonstrate similar perturbation responses, but also which components in the transcription processing pipeline define those shared responses.

## Discussion

Overall, this application of stochastic biophysical models to high-throughput genomics data demonstrates an alternative avenue for how we analyze, interpret, and develop hypotheses from large-scale perturbation. The *Monod* CME inference package and the meK-Means clustering algorithm, enable this analysis for noisy and discrete single-cell data, and extract physically interpretable parameters from the data as well as demonstrate methods that can be consistently extended to new measurements.

In this study, the biophysical models utilized in *Monod* and meK-Means focus on a relatively simple model of bursty transcription, and assume the effects of cell size are negligible. These methods additionally utilize CME models of biological systems where analytical solutions are available, solved through the *Monod* framework. However, recent developments in combining machine learning with biophysical models of transcription, for parameter inference [61, 62], suggest promising extensions of this work to simultaneously incorporate cell size effects on transcription and extend inference to more complex biophysical models, without analytical solutions, that can be simulated. Along these lines, though meK-Means does represent shared relationships between genes in order to cluster the cells, it does not learn more direct interactions between genes and how they may effect the biophysical parameters of the system. However, integration of such approaches with statistical and learning-based approaches for causal inference from perturbation data [8, 63, 64] would merge the learned interactions with their effects on transcriptional dynamics.

As is, the methods used in this study provide users a biophysical and statistical basis through which to interpret perturbation biology. For example, Chi-squared goodness-of-fit testing is used to reject genes which do not fit the assumed models of transcription [31]. To select between meK-Means clustering results under different values of the hyperparameter K, we can use criteria such as the Akaike Information Criterion (AIC) to determine which result better describes the data (and when increasing K does not improve fit) (Supplementary Fig. 4b,5b). In a similar vein, the Fisher information matrix is also calculated for the inferred parameters (see Methods) and can be used to assess their intrinsic uncertainty and information content (Supplementary Fig. 2), potentially aiding downstream, experimental design informed by biophysical properties of the biology [20].

This stochastic, biophysical approach to modeling large-scale, perturbation biology is also particularly promising given increasing focus on and development of experimental techniques to enhance the depth and breadth of capture of mRNA at different stages of processing [65, 66], which would greatly improve the quantity of genes that can be used for biophysical inference. Beyond mRNA, these physical models additionally emulate the kinds of inter-modality relationships present in other multimodal perturbation data. CME-based approaches have likewise demonstrated such physical modeling between mRNA and protein expression dynamics [29], and chromatin openness and mRNA transcription [67]. Thus, as perturbation genomics data become increasingly complex and multimodal, this work demonstrates a paradigm that aims for scalability not just in dataset size but also in interpretation, with methods that can extend to new measurements and modalities and provide physically interpretable insights into the cellular processes governing our molecular measurements.

## Methods

### A CME Model of Sequencing Data

The underlying biophysical model of the counts observed in the perturbation data, used for this analysis, describes bursty transcription of mRNA molecules, coupled to splicing and degradation of those molecules, followed by length-biased capture of the mRNA during the sequencing process which produces the final observed counts (Fig. 5) [28]. Given the sparse and discrete nature of scRNA-seq count data, we utilize the Chemical Master Equation (CME) to model these molecule counts as a function of the kinetic parameters of the biophysical model [29, 68]. The CME-formalism is particularly suited to discrete (low) count regimes and has been extensively used in the fluorescence transcriptomics literature to model the mRNA production and translation in single cells [29, 68–70].

The CME models the probability of molecule counts over time *t* i.e., *p*(*X, t*) where *X* is the random variable representing molecule counts or ‘states’ in the system. *p*(*X, t*) is defined through parameters *θ* which represent some rates of transitions between states [29, 68]. For the biophysical model described above our states are counts of unspliced and spliced mRNA represented by random variables *U, S*. The rates of transition are then defined as the transcription rate *k* (and burst size *b*), splicing at rate *β*, and degradation at rate *γ*. The molecules produced from the biological, transcription processes are then captured and sampled during the technical, sequencing process. The net capture rates for the unspliced and spliced molecules are defined through the *C^u^*and *λ^s^*parameters. The capture of unspliced mRNA is length-dependent (as greater length and thus internal poly(A) tracts can lead to more capture/binding events given a sequencing technology using a poly(T)-based system for mRNA capture), represented by *λ^u^*= *C^u^L_g_*, where *L_g_* represents the length of the gene *g*. All parameters ***θ*** = *{λ^u^, λ^s^, k, b, β, γ}* are modeled on a per-gene basis, i.e., *k_g_, b_g_, β_g_, γ_g_*. Together the parameters define how the probability of *U_g_, S_g_* counts evolves over time (Fig. 5).

This length-biased model of scRNA-seq data is developed and described in further detail in [28]. Here we fit the steady-state model to the datasets, when *t → ∞*, as we are looking at snapshot data of unsynchronized cells. In this regime, certain parameters are not uniquely identifiable. As described in [28], the splicing and degradation rates are defined relative to a transcription rate *k* (*β/k, γ/k*) where *k* = 1 without loss of generality. Likewise, the capture rate parameters represent a net capture rate encompassing Poisson length-biased poly(A) capture and binomial sampling during the sequencing and PCR steps (Fig. 5).

### Solving the CME Model with *Monod*

We infer or ‘fit’ parameters ***θ*** to the raw unspliced and spliced counts, which represent our observed molecule counts, by minimizing the KLD between the probability distribution induced by the CME model above, and the joint probability distribution (or histogram) of observed counts. This means, given the count matrices **U** and **S** *∈* R^N^*^×^*^G^ (for N cells and G genes), we optimize (per gene) for ***θ****_g_* = [*b_g_, β_g_/k_g_, γ_g_/k_g_, λ^u^, λ^s^*]:

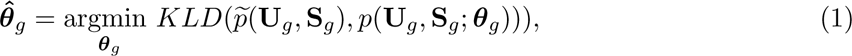

where *p* describes the observed histogram of counts [31].

To solve for the probability of molecule counts given the parameters of the biophysical model, we use the analytical solution derived in [29] that employs the generating function representation of our CME model. Thus, given a steady-state probability generating function (PGF) form, *H*, of *p*(*U_g_, S_g_*), we can solve for *p*(*U_g_, S_g_*; ***θ****_g_*):

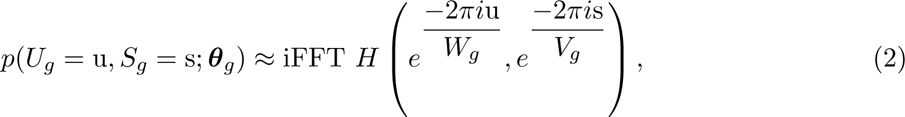

where u = 0*,., W_g_ −* 1, s = 0*,.., V_g_ −* 1 and *W_g_, V_g_* are sufficiently large, positive integers. This amounts to evaluating the PGF *H* around the complex unit circle and performing an inverse Fourier transform (denoted iFFT) to obtain the molecule count probabilities [29]. Here *W_g_* = max(**U***_g_*)*, V_g_* = max(**S***_g_*). The *Monod* package implements the numerical integration and optimization for the CME model [31].

The technical parameters *λ^u^, λ^s^*, which define the net capture rates described above, are fit using a grid search over potential capture rate values. The physical parameters are optimized at each grid point, using Equation 1. The ‘optimal’ technical parameters for a dataset are then determined as the capture rates at the grid point where the KLD across genes is lowest [31].

### The meK-Means Algorithm for Clustering

The meK-Means algorithm (Fig. 6) for clustering extends the model of the probability of molecule counts to include a latent variable *Z* that denotes the state (or cluster) a cell is in. Thus the probability of the counts is now *p*(*U, S, Z|****θ***) [33]. Here *Z* can take an (integer) value from 1 to the user-defined K. The posterior, *p*(*Z|U, S,* ***θ***), and parameters ***θ***, are unknown, so we take an Expectation Maximization (EM)-based approach to optimize the Q function (the objective function of our algorithm):

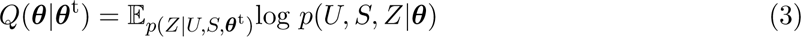

iterating between updating the posterior given parameter estimates ***θ***^t^ (E-step), and determining the MLE parameter estimates which then maximize *Q*(***θ****|****θ***^t^) (M-step). Here, t is the number of epochs.

The full algorithm details, benchmarking, and runtime metrics are described in [33]. Briefly, given the count matrices **U** and **S** *∈* R^N^*^×^*^G^ (for N cells and G genes), in the M-Step of the algorithm we use hard assignment of cells to a single latent state or cluster k, where the cell is assigned to a k such that

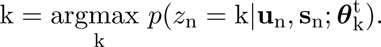

This is because the numerical procedure for obtaining parameter estimates for the defined CME model requires a histogram over observed counts (see Equation (1)). This mirrors the hard assignment of an observation to a cluster centroid in K-Means clustering (hence the ‘K-Means’ [71] in meK-Means).

In the E-Step, the posterior is updated as

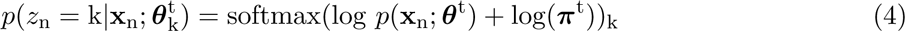

with *p*(**x**_n_; ***θ***^t^) as in Equation (2). ***π*** represents the mixing weights/priors of each cluster *Z* = k. The final cluster assignments are determined as **Z^^^** = [^z_1_*,…,* ^z_N_] and ^z_n_ = argmax *p*(*z*_n_ = k*|***x**_n_).

Though the user supplies a K to the meK-Means algorithm, since we are using hard-assignment of the cells to a cluster k, the algorithm can converge to fewer clusters (it is a more conservative approach than standard techniques such as K-Means or Leiden clustering).

### Dataset Preprocessing

For the CRISPRa, CRISPRi, drug combination NSC, and erlotinib-treated PC9 datasets, unspliced and spliced counts were generated with *kallisto|bustools* 0.26.0 [25, 72] from the FASTQ files. Specifically, the kb ref command with the--lamanno option was used to build exonic and intronic references for each genome. kb count with the--lamanno workflow and the -x 10xv2 or -x 10xv3 technology option was then used to obtain unspliced and spliced mRNA count matrices. Unspliced and spliced counts for the DEX-treated A549 cells were already provided in the original study.

The raw count matrices were used as input for *Monod* and meK-Means. The mm10 and GRCh38 (2020-A version) reference genomes were used for pseudoalignment, downloaded from 10x Genomics (https://support.10xgenomics.com/single-cell-gene-expression/software/downloads/latest). See Supplementary Table 1, for direct links to all data.

Cell barcodes with at least 10^4^ UMIs were selected for both CRISPR datasets, and greater than 1,500 UMIs for the NSC dataset. Cell barcodes annotated in the original study were used for the DEX-treated A549 cells. And the barcodes returned after the *kallisto|bustools* barcode filtering were used for the PC9 dataset.

To obtain the technical parameters for datasets where all samples were multiplexed and/or sequenced together, the CRISPR and NSC datasets, *Monod* inference was run on 3,000 genes with at least 0.01 average unspliced and spliced counts (and less than 400 maximum counts for each modality, per cell). 3,000 genes were selected per dataset, and we then selected conditions described in the text as well as all control conditions [2, 3, 27] and separately fit the genes for the conditions.

Optimal technical parameters from across the grid search (described above) were determined for each condition. Then for each dataset, the ‘consensus’ technical sampling parameters were set as the centroid of these optimal parameters across the conditions fit. For the A549 DEX and PC9 erlotinib datasets, 500 genes were selected by the same filter, and physical parameters inferred using the same technical parameter grid for each condition in these datasets separately. Since the conditions within these datasets were not sequenced together, the optimal technical parameters for each condition were used for downstream analysis.

### *Monod* Inference for Condition-Specific Kinetics

For the A549 DEX and CRISPRa conditions, 3,000 genes (in the CRISPRa case the same genes as above) were fit with *Monod* under the bursty model of transcription with length-biased sampling as described above. For the CRISPRa data we additionally ensured that the target genes corresponding to each intervention condition were included.

Parameter fold changes (FCs) as shown in Fig. 2e, were computed against the parameters inferred using the cells from all control guide conditions combined (following the analysis in the study, and only combing the control guides not rejected by the original study [2]). Standard deviation error bars in Fig. 2e (right), are thus calculated given the FCs of the condition versus each control condition individually.

### Nuclear Speckle Proximity Analysis

To assess the genomic location and speckle proximity of the genes used for inference from the CRISPRa data, we obtained the chromosomal coordinates of each gene from Ensembl BioMart release 112 (GRCh38.p14), for the genes displaying increased fold changes in the perturbation condition. ‘Speckle proximity’ scores were obtained from the authors of [45], where the SPRITE method is used to capture multi-way interactions between DNA (and RNA) molecules in the nucleus. Proximity to nuclear speckles (nuclear bodies with mRNA splicing and transcription machinery) was determined by the frequency of contacts/interactions between 1Mb regions of the genome to regions determined as transcriptionally ‘active’ hubs (where active hubs corresponded to organization around nuclear speckles) [45]. Regions considered speckle ‘close’ were denoted as loci with proximity scores/frequencies greater than the 95th percentile (across the genome) [45]. To plot genes along the genome we used the shinyCircos web application [73].

### Marker Gene Determination and Parameter Uncertainty

To define differentially expressed genes at the parameter level, DE-*θ* genes, we looked for genes displaying log_2_FCs (log_2_ fold changes) greater than 2 in magnitude between the parameters of the two conditions or populations being compared. Genes that were not DE at the spliced mean-level, were defined as having log_2_FCs (between average expression values) of magnitude less than 1.0. We did not consider rejected genes (genes with poor model fits) for DE analysis. Rejected genes were determined by (1) genes with inferred parameters that remained close to the initialized parameter bounds, i.e., where gradient descent did not produce reliable estimates [31], (2) *<* 0.05*/*G p-values obtained from Chi-squared tests between observed count histograms and the distributions induced by the inferred parameters (after Bonferroni correction for the G genes tested), and (3) Hellinger distances *>* 0.05 between these two distributions [31].

For parameter estimates, standard error, *σ*, values can be calculated from the square root of the diagonals of the inverse Fisher Information Matrix (FIM). The FIM is calculated as the Hessian matrix of the KLDs between the observed count histograms and the distributions induced by the inferred parameters. Analysis of the FIM for parameters is shown in Supplementary Fig. 2, to assess how informative the data is regarding the biophysical parameters of interest.

### Parameter Prediction Models

#### Definition of Predictive Models

For the predictive models of kinetics in combined perturbation conditions, we focused on additive and multiplicative models of transcript production to describe the changes in *b* or *k*. As described in [23, 47], multiplicative changes at both the level of mean expression FC and transcription/forward rate can result from the ‘one-step recruitment’ model [47] of transcription regulation, where the combined interventions behave in an additive manner to alter the free energy of the system, resulting in multiplicative effects at the level of the rates and observed expression FCs.

For this study, the multiplicative models of the parameters in the combined conditions (relative to control) are then:

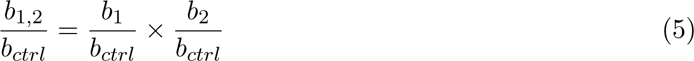

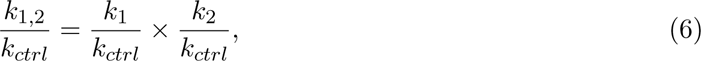

where 1 and 2 denote the single-perturbation conditions, and ‘1,2’ denotes the combined condition. Parameters denoted with ‘ctrl’ represent the control condition’s parameters. These models also assume that changes in *β/k* represent changes in *k*, i.e., *β/k* and *γ/k* change together. Given that our biophysical model does not have a separate *k* parameter, we model these changes through *β/k* (denoted below as *β^′^* for convenience). Thus we rewrite Equation 6 as:

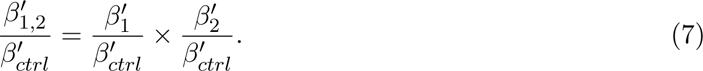

As also described in [23, 47], additive effects are also observed at the level of expression and can be derived from independent effects of the perturbations to, in parallel, catalyze or reduce the forward rate [47].

For this study, the additive models are then:

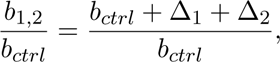

where *b*_1_ = *b_ctrl_* + Δ_1_ and *b*_2_ = *b_ctrl_* + Δ_2_. This can be rewritten as:

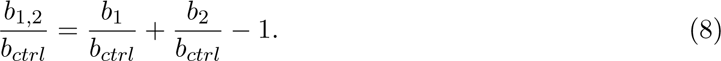

Likewise for *k*:

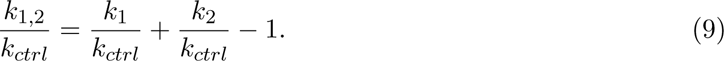

To obtain a formulation in terms of *β^′^*:

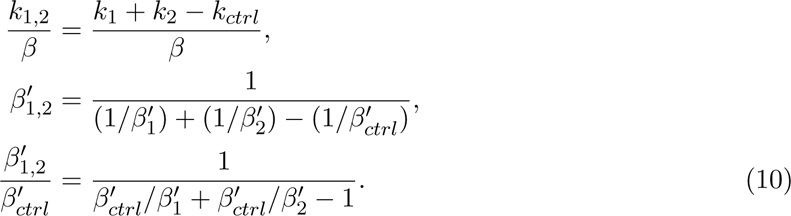

The same models can be derived from multiplicative and additive changes observed at the level of mean expression, if for example, either *b* or *k* is driving those moments’ changes, based on the first moment derivations for the bursty model of transcription. For reference, the first moments of the length-biased model of transcription are:

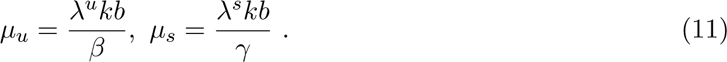

Currently, these predictive models do not incorporate multiple steps in transcription production [22] as this uses a simplified version of the two-state telegraph model, where in the bursty-limit there is one forward rate *k* or *k_on_* (inactive to active state transition) and the burst size *b* represents a ratio between the *k_ini_* rate (rate of production) and *k_off_* rate (active to inactive transition), as *k_ini_, k_off_ → ∞* [28].

### Gene Selection for Prediction

As described in the CRISPRa data study [2], a random forest regression model was used to select the top genes (from the 3,000 selected above) which best separated the single-guide (single-perturbation) conditions from the control conditions (and each other). Unlike [2], we did not include the third, combined condition in this selection/model. We additionally tested a simpler logistic regression classifier as well. Given results in Supplementary Fig. 3, it is possible that as suggested in [2], the random forest model is able to better detect genes with distributional changes in expression, not necessarily discernible through FC, but that are more amenable to prediction. The sklearn functions ExtraTreesClassifier or LogisticRegression were used for each regression model, with the top-weighted 100 genes used for prediction.

For the CRISPRi and NSC datasets, 3,000 genes were fit with *Monod* under the bursty model of transcription with length-biased sampling as described above. The same gene selection procedures for prediction as used for the CRISPRa data were applied to these datasets. For parameter FC comparisons we used the same pairing of control guides and combined perturbation conditions as in the CRISPRi study [27], and used the high EGF (no RA, no BMP4) conditions for the NSC controls [3].

To aid interpretation and analysis of additive/multiplicative changes, we then selected for genes in the CRISPRa parameters where single-conditions acted in the same ‘direction’ (activation or repression) and where *k* was likely affected, i.e., where *β/k* and *γ/k* FCs were in the same direction as each other within and between the two single-perturbation conditions. We then selected repressed genes, where the average expression log_2_FC was *< −*0.5 for unspliced counts (i.e., production was decreased).

### Assigning Representative Predictive Models

To determine which predictive model best represented the observed behavior in the combined condition, we looked to see if the predicted FC per parameter, of each model, fell within the 95% C.I. of the observed parameter FC in the combined condition (constructed from the standard deviation of the FCs calculated in comparison to the individual control conditions) [23]. When only one model fell in that range, the parameter’s behavior was denoted as that model (‘Multiplicative’ or ‘Additive’). If both models fell in that range, we denoted those parameters’ behaviors as ‘Ambiguous’. If the observed parameter FC range was greater in magnitude than all predictions, this was denoted as ‘Super Multiplicative’, and likewise if the observed FC range was smaller in magnitude than all model predictions, this was ‘Sub Additive’. We assigned predictive models in this way for the transcription rate *k*, for the CRISPRa conditions (above) where it was likely *k* was altered in both single perturbations.

### HOMER Analysis of Binding Motifs

To determine what genomic features might be shared among genes displaying similar combinatorial strategies/predictive models, the HOMER software [51] for motif discovery was used to detect transcription factor binding domains in the genes’ promoter regions. We used the HOMER workflow for discovery in promoter regions, specifically findMotifs.pl, to determine the presence of ‘known motifs’. Results and command line arguments are in the code repository.

### meK-Means Clustering Analysis

For the DEX-treated A549 cells dataset, the gene list of potential cortisol-responsive genes from [35] was used for inference. For the Day 3 cells in the erlotinib-treated PC9 cells study, 2,000 HVGs were selected using the scanpy [56] highly variable genes using the spliced matrix, as is standard. Additionally genes discussed throughout the original study [59] (as previously or newly described markers of resistance) were also included. For both datasets, these lists of genes were then filtered for genes with dispersion *≥* 1.5, minimum unspliced and spliced average counts of 0.02, and less than a 50x ratio between spliced to unspliced average counts [33]. This is a more stringent filtering criteria than for the analysis above, since genes are no longer treated completely independently in this clustering task (the genes’ fits impact each other), as opposed to fitting parameters with the standard *Monod* pipeline for each gene in parallel.

For the comparison to Leiden clustering, the input data matrices underwent standard normalization procedures in scanpy [56] (read-depth normalization and log1p transformation). We then used the kneighbors graph function from sklearn to create a nearest neighors graph of these processed matrices as input into the Leiden algorithm. We used a default 30 neighbors and resolution of 0.1, following standard scanpy procedure. To compare the overlapping nature of the clustering results from each of the input matrix options, we calculated the Adjusted Rand Index (ARI) score between all pairs of Leiden cluster results. For the ARI score, 1.0 denotes overlapping assignments and 0.0 represents poor or random assignments.

For the meK-Means runs, we tested K values of K=2,5,10 to cover a range of possibilities. We then used the Q function’s behavior (objective function in The meK-Means Algorithm for Clustering) and the Akaike Information Criterion (AIC) to compare model fits over a range of K values (see Supplementary Fig. 4b,5b). The AIC here is defined as:

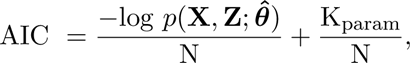

where K_param_ = 3 *×* G *×* K_Final_ + (K_Final_ *−* 1), for G genes and N cells, and where K_Final_ is the final number of clusters k that are assigned to cells. The AIC thus incorporates model fit through the likelihood of the data, and a penalty for the complexity of the model, through the number of parameters.

Using *Monod* (without meK-Means clustering), we separately fit gene parameters for the same genes as fit using meK-Means on the Day 3 PC9 cells and Day 0 cells separately, for comparison (see Supplementary Fig. 6).

All *Monod* and meK-Means analyses were run on an Intel Xeon Gold 6152 at 2.10 GHz using up to 30 cores.

## Data and Code Availability

Links to the original data used in this study are provided in Supplementary Table 1.

All code used to generate the figures and results in the paper is available at https://github.com/pachterlab/CGP_2024_2. A meK-Means [33] tutorial can be found here https://github.com/pachterlab/CGP_2023/blob/main/example_meKMeans_notebook.ipynb. The algorithm is implemented as a part of the *Monod* package [31] for single-cell, CME-based parameter inference, whose documentation can be found here https://monod-examples.readthedocs.io/en/latest/.

## Acknowledgements

We thank Thomas Norman and Joseph Replogle for providing the differentially expressed genes used in the CRISPR intervention studies, Prashant Bhat for providing the SPRITE proximity data, and Meichen Fang for helpful feedback on the manuscript. This work was partially funded by NIH 5UM1HG012077-03.

## References

1. Dixit, A. et al. Perturb-Seq: Dissecting Molecular Circuits with Scalable Single-Cell RNA Profiling of Pooled Genetic Screens. en. Cell 167, 1853–1866.e17 (Dec. 2016).

2. Norman, T. M. et al. Exploring genetic interaction manifolds constructed from rich single-cell phenotypes. Science (Aug. 2019).

3. Gehring, J., Hwee Park, J., Chen, S., Thomson, M. & Pachter, L. Highly multiplexed single-cell RNA-seq by DNA oligonucleotide tagging of cellular proteins. Nat. Biotechnol. 38, 35–38 (Jan. 2020).

4. Chen, S. et al. Dissecting heterogeneous cell populations across drug and disease conditions with PopAlign. Proc. Natl. Acad. Sci. U. S. A. 117, 28784–28794 (Nov. 2020).

5. Inecik, K., Uhlmann, A., Lotfollahi, M. & Theis, F. MultiCPA: Multimodal Compositional Perturbation Autoencoder en. July 2022.

6. Frangieh, C. J. et al. Multimodal pooled Perturb-CITE-seq screens in patient models define mechanisms of cancer immune evasion. Nat. Genet. 53, 332–341 (Mar. 2021).

7. Freimer, J. W. et al. Systematic discovery and perturbation of regulatory genes in human T cells reveals the architecture of immune networks. Nat. Genet. 54, 1133–1144 (Aug. 2022).

8. Chevalley, M., Roohani, Y., Mehrjou, A., Leskovec, J. & Schwab, P. CausalBench: A Large-scale Benchmark for Network Inference from Single-cell Perturbation Data. arXiv: 2210.17283 [cs.LG] (Oct. 2022).

9. Sullivan, D. K. et al. kallisto, bustools, and kb-python for quantifying bulk, single-cell, and single-nucleus RNA-seq. en. bioRxiv (Jan. 2024).

10. Chamberlin, J. T., Lee, Y., Marth, G. T. & Quinlan, A. R. Differences in molecular sampling and data processing explain variation among single-cell and single-nucleus RNA-seq experiments en. July 2023.

11. Mayere, C., et al. Single-cell transcriptomics reveal temporal dynamics of critical regulators of germ cell fate during mouse sex determination. en. FASEB J. 35, e21452 (Apr. 2021).

12. Wang, B.-D. & Lee, N. H. Aberrant RNA Splicing in Cancer and Drug Resistance. Cancers 10 (Nov. 2018).

13. Conboy, J. G. Developmental regulation of RNA processing by Rbfox proteins. Wiley Interdiscip. Rev. RNA 8 (Mar. 2017).

14. Lotfollahi, M., Wolf, F. A. & Theis, F. J. scGen predicts single-cell perturbation responses. Nat. Methods 16, 715–721 (Aug. 2019).

15. Jiang, J. et al. D-SPIN constructs gene regulatory network models from multiplexed scRNA-seq data revealing organizing principles of cellular perturbation response. bioRxiv (May 2023).

16. Ji, Y., Lotfollahi, M., Wolf, F. A. & Theis, F. J. Machine learning for perturbational single-cell omics. Cell Syst 12, 522–537 (June 2021).

17. Chari, T. & Pachter, L. The specious art of single-cell genomics. PLoS Comput. Biol. 19, e1011288 (Aug. 2023).

18. Ahlmann-Eltze, C. & Huber, W. Comparison of transformations for single-cell RNA-seq data. en. Nat. Methods 20, 665–672 (May 2023).

19. Tyler, S. R., Bunyavanich, S. & Schadt, E. E. PMD Uncovers Widespread Cell-State Erasure by scRNAseq Batch Correction Methods en. Nov. 2021.

20. Fox, Z. R., Neuert, G. & Munsky, B. Optimal Design of Single-Cell Experiments within Temporally Fluctuating Environments. Complexity 2020 (June 2020).

21. Yuan, B. et al. CellBox: Interpretable Machine Learning for Perturbation Biology with Application to the Design of Cancer Combination Therapy. Cell Systems 12, 128–140.e4 (Feb. 2021).

22. Bartman, C. R. et al. Transcriptional Burst Initiation and Polymerase Pause Release Are Key Control Points of Transcriptional Regulation. en. Mol. Cell 73, 519–532.e4 (Feb. 2019).

23. Sanford, E. M., Emert, B. L., Cote, A. & Raj, A. Gene regulation gravitates toward either addition or multiplication when combining the effects of two signals. en. Elife 9 (Dec. 2020).

24. Hjorleifsson, K. E., Sullivan, D. K., Holley, G., Melsted, P. & Pachter, L. Accurate quantification of single-nucleus and single-cell RNA-seq transcripts en. Dec. 2022.

25. Melsted, P. et al. Modular and efficient pre-processing of single-cell RNA-seq en. July 2019.

26. Cao, J., Zhou, W., Steemers, F., Trapnell, C. & Shendure, J. Sci-fate characterizes the dynamics of gene expression in single cells. Nat. Biotechnol. 38, 980–988 (Aug. 2020).

27. Replogle, J. M. et al. Combinatorial single-cell CRISPR screens by direct guide RNA capture and targeted sequencing. Nat. Biotechnol. 38, 954–961 (Aug. 2020).

28. Gorin, G. & Pachter, L. Length biases in single-cell RNA sequencing of pre-mRNA. Biophys Rep (N Y) 3, 100097 (Mar. 2023).

29. Bokes, P., King, J. R., Wood, A. T. A. & Loose, M. Exact and approximate distributions of protein and mRNA levels in the low-copy regime of gene expression. J. Math. Biol. 64, 829–854 (Apr. 2012).

30. Cai, L., Friedman, N. & Xie, X. S. Stochastic protein expression in individual cells at the single molecule level. Nature 440, 358–362 (Mar. 2006).

31. Gorin, G. & Pachter, L. Monod: mechanistic analysis of single-cell RNA sequencing count data en. June 2022.

32. Burkhardt, D. B. et al. Quantifying the effect of experimental perturbations at single-cell resolution. Nat. Biotechnol. 39, 619–629 (May 2021).

33. Chari, T., Gorin, G. & Pachter, L. Biophysically Interpretable Inference of Cell Types from Multimodal Sequencing Data. bioRxiv (Sept. 2023).

34. Anders, S. & Huber, W. Differential expression analysis for sequence count data. Genome Biol. 11, R106 (Oct. 2010).

35. Reddy, T. E. et al. Genomic determination of the glucocorticoid response reveals unexpected mechanisms of gene regulation. Genome Res. 19, 2163–2171 (Dec. 2009).

36. Kurihara-Shimomura, M. et al. Zinc finger AN1-type containing 4 is a novel marker for predicting metastasis and poor prognosis in oral squamous cell carcinoma. J. Clin. Pathol. 71, 436–441 (May 2018).

37. Pudil, R. et al. Plasma glycogen phosphorylase BB is associated with pulmonary artery wedge pressure and left ventricle mass index in patients with hypertrophic cardiomyopathy. Clin. Chem. Lab. Med. 48, 1193–1195 (Aug. 2010).

38. Syx, D. et al. The RIN2 syndrome: a new autosomal recessive connective tissue disorder caused by deficiency of Ras and Rab interactor 2 (RIN2). en. Hum. Genet. 128, 79–88 (July 2010).

39. Perez-Pinera, P. et al. Synergistic and tunable human gene activation by combinations of synthetic transcription factors. Nat. Methods 10, 239–242 (Mar. 2013).

40. Liang, Y. et al. Gene activation guided by nascent RNA-bound transcription factors. Nat. Commun. 13, 7329 (Nov. 2022).

41. Nicolas, D., Zoller, B., Suter, D. M. & Naef, F. Modulation of transcriptional burst frequency by histone acetylation. en. Proc. Natl. Acad. Sci. U. S. A. 115, 7153–7158 (July 2018).

42. Dar, R. D. et al. Transcriptional burst frequency and burst size are equally modulated across the human genome. Proc. Natl. Acad. Sci. U. S. A. 109, 17454–17459 (Oct. 2012).

43. Wang, Y., Qi, J., Shao, J. & Tang, X.-Q. Signaling Mechanism of Transcriptional Bursting: A Technical Resolution-Independent Study. en. Biology 9 (Oct. 2020).

44. Li, Y. et al. MiR-218 Inhibits Erythroid Differentiation and Alters Iron Metabolism by Targeting ALAS2 in K562 Cells. en. Int. J. Mol. Sci. 16, 28156–28168 (Nov. 2015).

45. Quinodoz, S. A. et al. Higher-Order Inter-chromosomal Hubs Shape 3D Genome Organization in the Nucleus. Cell 174, 744–757.e24 (July 2018).

46. Lotfollahi, M. et al. Predicting cellular responses to complex perturbations in high-throughput screens. Mol. Syst. Biol., e11517 (May 2023).

47. Scholes, C., DePace, A. H. & Śanchez, A. Combinatorial Gene Regulation through Kinetic Control of the Transcription Cycle. Cell Syst 4, 97–108.e9 (Jan. 2017).

48. Sears, T. K. & Angelastro, J. M. The transcription factor ATF5: role in cellular differentiation, stress responses, and cancer. Oncotarget 8, 84595–84609 (Oct. 2017).

49. Huang, C., Zhan, J.-F., Chen, Y.-X., Xu, C.-Y. & Chen, Y. LncRNA-SNHG29 inhibits vascular smooth muscle cell calcification by downregulating miR-200b-3p to activate the *α*-Klotho/FGFR1/FGF23 axis. Cytokine 136, 155243 (Dec. 2020).

50. Hu, C. et al. Effects of miR-210-3p on the erythroid differentiation of K562 cells under hypoxia. Mol. Med. Rep. 24 (Aug. 2021).

51. Heinz, S. et al. Simple combinations of lineage-determining transcription factors prime cisregulatory elements required for macrophage and B cell identities. en. Mol. Cell 38, 576–589 (May 2010).

52. Yang, S.-H., Bumpass, D. C., Perkins, N. D. & Sharrocks, A. D. The ETS domain transcription factor Elk-1 contains a novel class of repression domain. en. Mol. Cell. Biol. 22, 5036–5046 (July 2002).

53. Traag, V. A., Waltman, L. & van Eck, N. J. From Louvain to Leiden: guaranteeing well-connected communities. Sci. Rep. 9, 5233 (Mar. 2019).

54. Hao, Y. et al. Integrated analysis of multimodal single-cell data. Cell (May 2021).

55. Lin, X., Tian, T., Wei, Z. & Hakonarson, H. Clustering of single-cell multi-omics data with a multimodal deep learning method. Nat. Commun. 13, 7705 (Dec. 2022).

56. Wolf, F. A., Angerer, P. & Theis, F. J. SCANPY: large-scale single-cell gene expression data analysis. en. Genome Biol. 19, 15 (Feb. 2018).

57. Heumos, L. et al. Best practices for single-cell analysis across modalities. Nat. Rev. Genet. 24, 550–572 (Aug. 2023).

58. Cao, Z.-J. & Gao, G. Multi-omics single-cell data integration and regulatory inference with graph-linked embedding. Nat. Biotechnol. 40, 1458–1466 (Oct. 2022).

59. Aissa, A. F. et al. Single-cell transcriptional changes associated with drug tolerance and response to combination therapies in cancer. en. Nat. Commun. 12, 1628 (Mar. 2021).

60. Shang, J., Zhang, X., Hou, G. & Qi, Y. HMMR potential as a diagnostic and prognostic biomarker of cancer—speculation based on a pan-cancer analysis. Frontiers in Surgery 9 (2023).

61. Carilli, M., Gorin, G., Choi, Y., Chari, T. & Pachter, L. Biophysical modeling with variational autoencoders for bimodal, single-cell RNA sequencing data. bioRxiv (May 2023).

62. Sukys, A., Ocal, K. & Grima, R. Approximating solutions of the Chemical Master equation using neural networks. iScience 25, 105010 (Sept. 2022).

63. Squires, C., Seigal, A., Bhate, S. & Uhler, C. Linear Causal Disentanglement via Interventions. arXiv: 2211.16467 [stat.ML] (Nov. 2022).

64. Lopez, R. et al. Learning Causal Representations of Single Cells via Sparse Mechanism Shift Modeling in Proceedings of the Second Conference on Causal Learning and Reasoning (eds van der Schaar, M., Zhang, C. & Janzing, D.) 213 (PMLR, 2023), 662–691.

65. Xiong, Y. et al. A Comparison of mRNA Sequencing with Random Primed and 3’-Directed Libraries. Sci. Rep. 7, 14626 (Nov. 2017).

66. Amarasinghe, S. L. et al. Opportunities and challenges in long-read sequencing data analysis. Genome Biol. 21, 30 (Feb. 2020).

67. Felce, C., Gorin, G. & Pachter, L. A Biophysical Model for ATAC-seq Data Analysis en. Jan. 2024.

68. Singh, A. & Bokes, P. Consequences of mRNA transport on stochastic variability in protein levels. en. Biophys. J. 103, 1087–1096 (Sept. 2012).

69. Munsky, B. & Khammash, M. The finite state projection algorithm for the solution of the chemical master equation. J. Chem. Phys. 124, 044104 (Jan. 2006).

70. Munsky, B., Fox, Z. & Neuert, G. Integrating single-molecule experiments and discrete stochastic models to understand heterogeneous gene transcription dynamics. Methods 85, 12–21 (Sept. 2015).

71. MacQueen, J. et al. Some methods for classification and analysis of multivariate observations in Proceedings of the fifth Berkeley symposium on mathematical statistics and probability 1> (books.google.com, 1967), 281–297.

72. Bray, N. L., Pimentel, H., Melsted, P. & Pachter, L. Erratum: Near-optimal probabilistic RNA-seq quantification. Nat. Biotechnol. 34, 888 (Aug. 2016).

73. Wang, Y. et al. shinyCircos-V2.0: Leveraging the creation of Circos plot with enhanced usability and advanced features. Imeta 2, e109 (May 2023).

