## Supplementary Materials for "Stochastic Modeling of Biophysical Responses to Perturbation"

### Stochastic Modeling of Biophysical Responses to Perturbation
#### Supplementary Materials

Tara Chari[1], Gennady Gorin[2], and Lior Pachter[1,3]

[1]Division of Biology and Biological Engineering,
California Institute of Technology, Pasadena, California

[2]Fauna Bio, Emeryville, California

[3]Department of Computing and Mathematical Sciences,
California Institute of Technology, Pasadena, California

July 4, 2024

### Tables

#### Datasets

All datasets used in this study are listed in Supplementary Table 1, with links to the original data.

| Dataset | Technology | FASTQs | Loom (U/S) |
|---|---|---|---|
| DEX-treated A549 Cells | sci-fate | GSE131351 | shendure-web.gs.washington.edu |
| CRISPRa Combinatorial Perturb-seq | 10xv2 | GSE133344 | 10.22002/ahjyk-gsj16 |
| CRISPRi Combinatorial Perturb-seq | 10xv3 | GSE146194 | 10.22002/tmx6v-57n22 |
| Drug-Combo Mouse NSCs | 10xv2 | 10.22002/D1.1311 | 10.22002/rt43q-s5v58 |
| Erlotinib-treated PC9s | 10xv3 | GSE148465 | 10.22002/cyr5a-ws203 |

**Supplementary Table 1. Dataset Metadata.** Datasets used for all analyses. GEO accession values or DOIs provided for FASTQs and Loom files (containing U and S counts).

### Figures

#### Kinetic Effects of Perturbation on Transcription

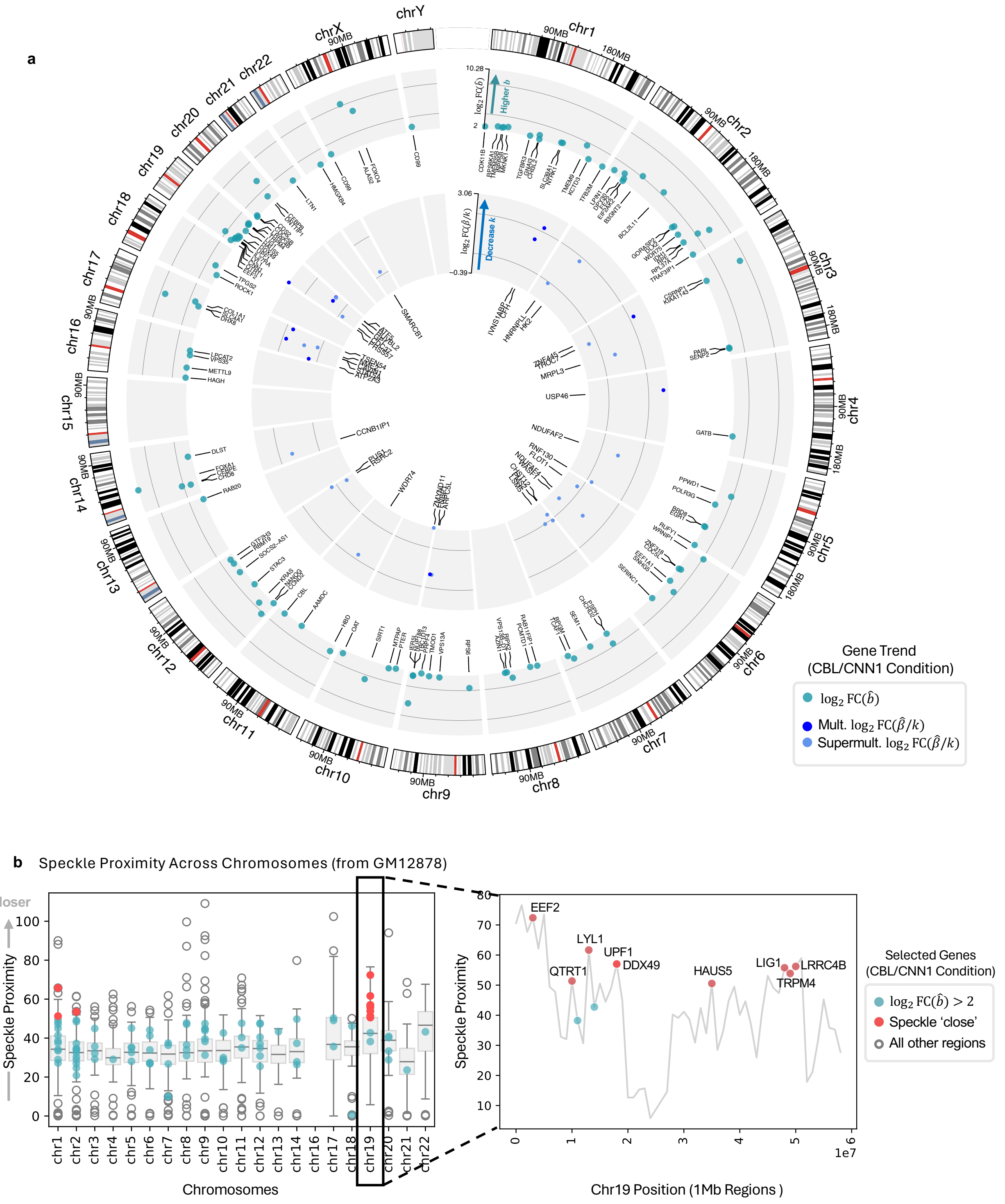

**Supplementary Fig 1. Shared Genome Localization and Kinetics. a** Circos [73] plot of genes with increased burst size (outer ring) or repressed transcription rate (inner ring) in the *CBL/CNN1* dual guide CRISPRa condition. Repressed genes denoted through multiplicative or super-multiplicative combination of individual perturbations. **b** (Left) Speckle proximity scores from [45] shown for genes from **a** with increased burst size, with respect to all other loci. Red dots denote genes with increased burst size *and* proximity scores in the 95th percentile (denoted as speckle 'close'). Proximity scores from GM12878 lymphoblastoid cells [45].

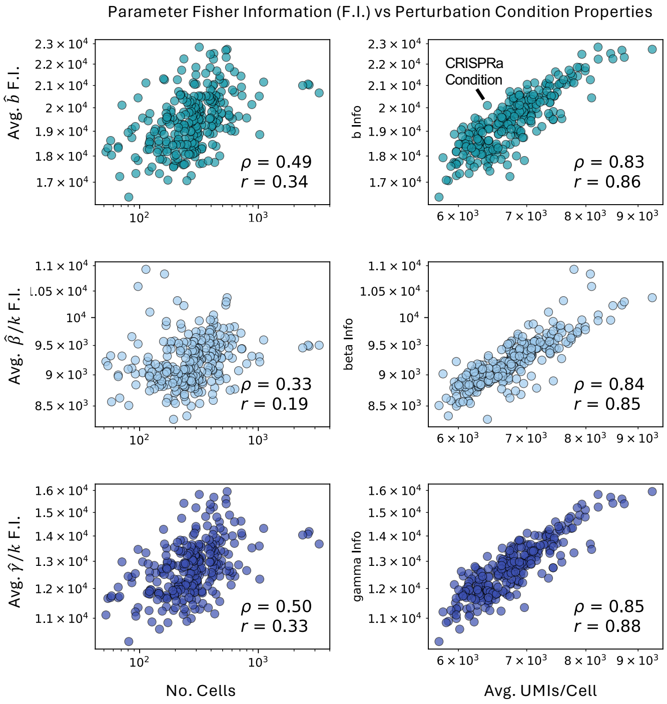

**Supplementary Fig 2. Fisher Information Content of CRISPRa Perturbations.** Average Fisher Information (F.I.) calculated per parameters for each CRISPRa condition. F.I. plotted against the number of cells in the condition or average UMIs per cell in the condition.(see Methods). Spearman and Pearson correlation are denoted by $\rho$ and $r$ respectively.

### Predictive Models of Combinatorial Perturbations

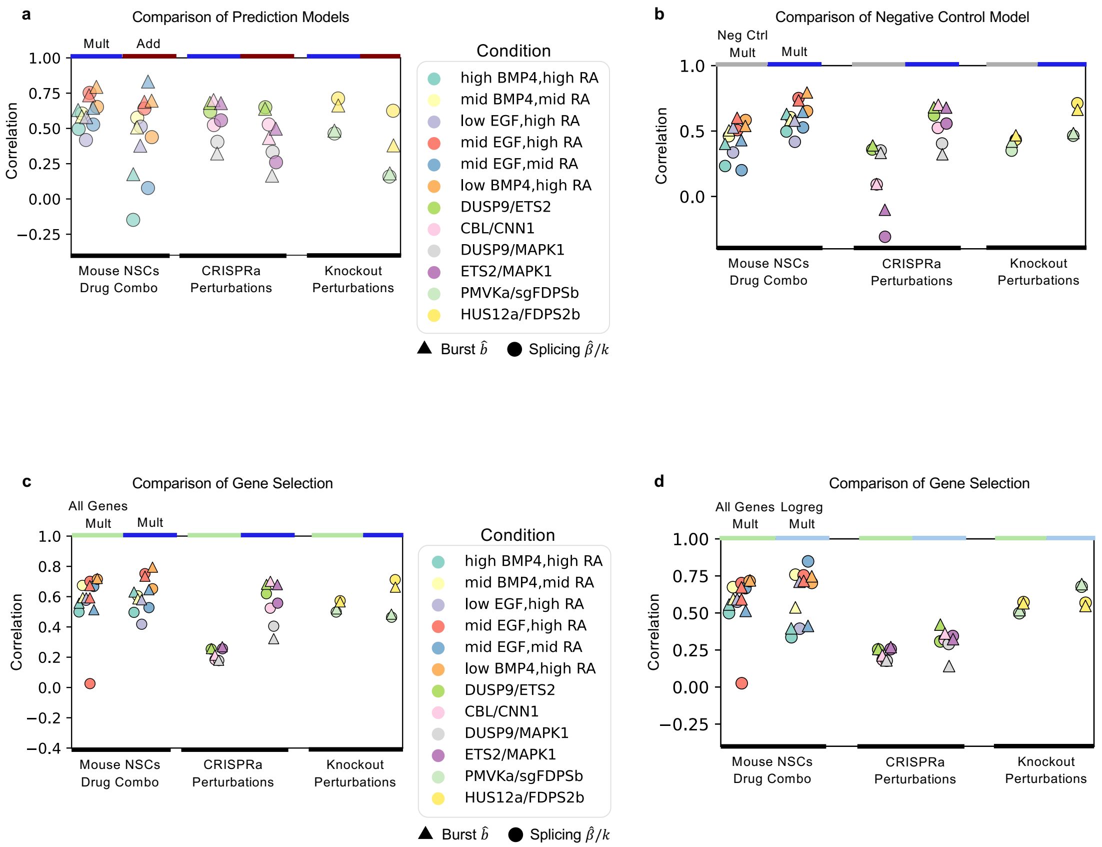

**Supplementary Fig 3. Predictive Model Comparisons Across Datasets. a** Pearson correlation of predicted parameters under the 'Mult' or 'Add' model for three datasets, across dual perturbation conditions in the datasets. Genes selected as done in [2], see Methods. Predictions shown for burst size and splicing rate. **b** Pearson correlation of predicted parameters under the 'Mult' model compared to the 'Neg Ctrl Mult' model (see Methods), across the same datasets and conditions. **c** Pearson correlation of predicted parameters under the 'Mult' model compared to the 'All Genes Mult' model, i.e., the same model applied to all genes without filtering/selection (see Methods), across the same datasets and conditions. **d** Pearson correlation of predicted parameters under the 'All Genes Mult' model compared to the 'Logreg Mult' model, i.e., the 'Mult' model applied to genes selected by logistic regression (see Methods).

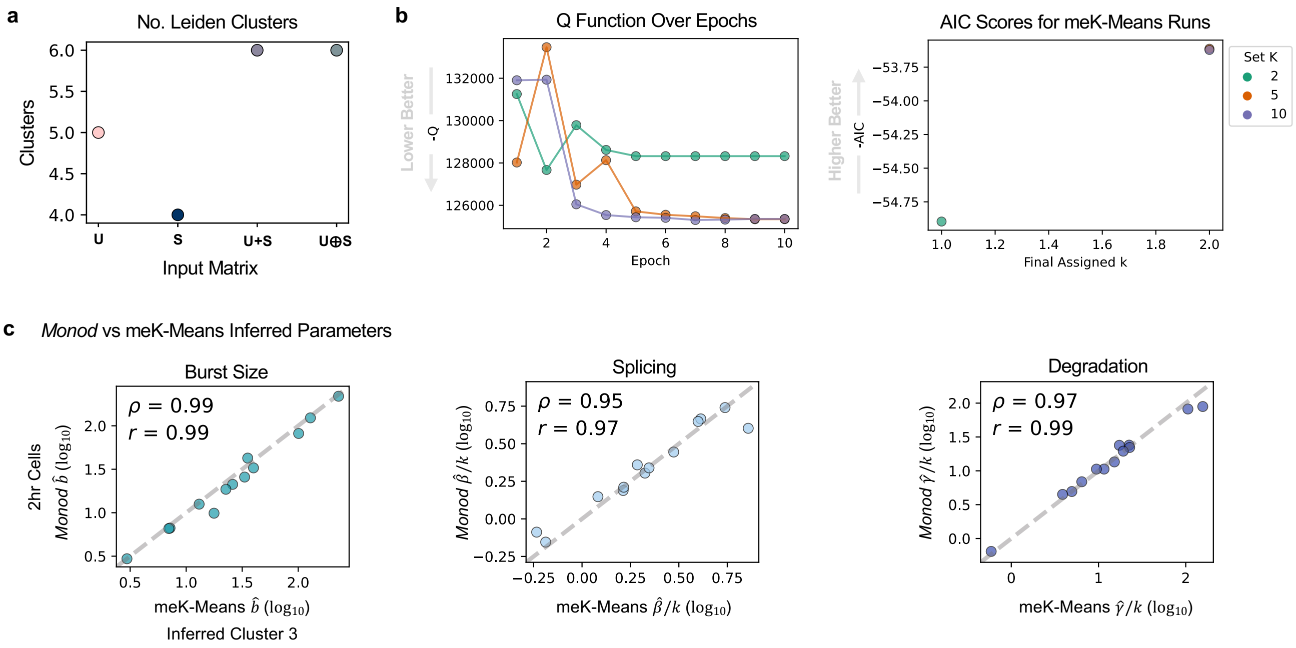

**Supplementary Fig 4. Clustering of DEX-Treated Cells. a** Number of clusters determined by Leiden clustering for all input matrix options. **b** -Q function and –AIC scores shown for meK-Means runs with K=2, 5, or 10 (see Methods). **c** Correlation of inferred parameters from *Monod* versus meK-Means inferred parameters (for genes used in clustering). Spearman and Pearson correlation are denoted by $\rho$ and $r$ respectively.

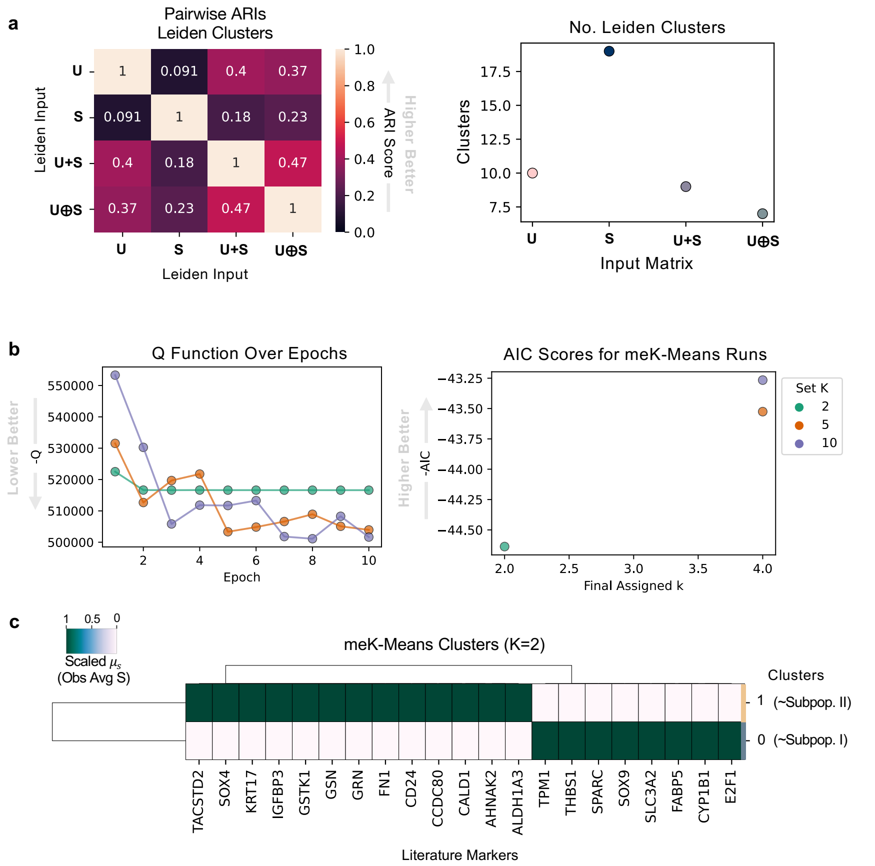

**Supplementary Fig 5. Clustering of Erlotinib-Resistant Cells. a** (Left) Pairwise ARI scores between Leiden clustering results across input matrix options. (Right) Number of clusters from Leiden results, across matrix input options. textbfb -Q function and –AIC scores shown for meK-Means runs with K=2, 5, or 10 (see Methods). **c** Hierarchical dendrogram plot of meK-Means inferred clusters based on average gene expression (scaled down the columns) of literature (published) markers. Clusters generally correspond to the Subpopulations I and II in [59].

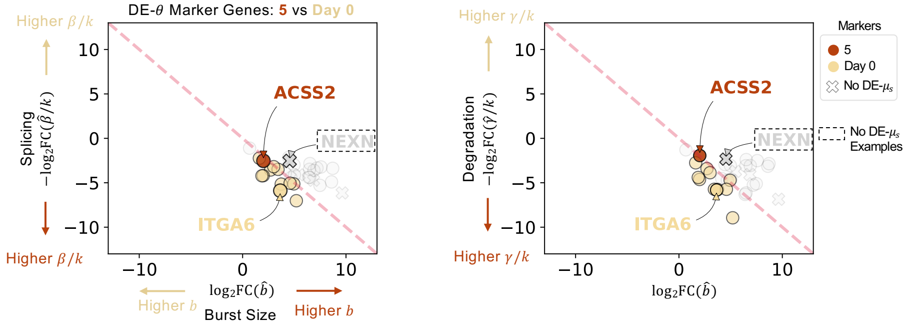

**Supplementary Fig 6. DE-$\theta$ Analysis Between DTP and Day 0 Cells.** 'DE' or 'Differentially-Expressed' genes at the parameter level ($\theta$) shown between the inferred cluster 5 cells and Day 0 cells. Genes in dashed box denote genes where DE is detected at the parameter level but not at the observed, mean S-level. (Left) Splicing rate vs burst size shown for genes. (Right) Degradation rate versus burst size shown. Grey genes denote ambiguous markers, or non-significant FCs. *NEXN* denotes an ambiguous gene with significant parameter FCs.
